# Host adaptation drives genetic diversity in a vector-borne disease system

**DOI:** 10.1101/2022.05.19.492734

**Authors:** Matthew A. Combs, Danielle M. Tufts, Ben Adams, Yi-Pin Lin, Sergios-Orestis Kolokotronis, Maria A. Diuk-Wasser

## Abstract

The range of hosts a pathogen can infect is a key trait influencing human disease risk and reservoir host infection dynamics. *Borrelia burgdorferi* sensu stricto (*Bb*), an emerging zoonotic pathogen, causes Lyme disease and is widely considered a host generalist, commonly infecting mammals and birds. Yet the extent of intraspecific variation in *Bb* host breadth, its role in determining host competence and potential implications to human infection remain unclear. We conducted a long-term study of *Bb* diversity, defined by the polymorphic *ospC* locus, across white-footed mice, passerine birds, and tick vectors leveraging long-read amplicon sequencing. Our results reveal strong variation in host breadth across *Bb* genotypes, exposing a spectrum of genotype-specific host-adapted phenotypes. We found support for multiple niche polymorphism maintaining *Bb* diversity in nature and little evidence of temporal shifts in genotype dominance, as would be expected under negative frequency-dependent selection. Passerine birds support the circulation of several human invasive strains in the local tick population and harbor greater *Bb* genotypic diversity compared to white-footed mice. Mouse-adapted *Bb* genotypes exhibited longer persistence in individual mice compared to non-adapted genotypes and infection communities infecting individual mice preferentially became dominated by mouse-adapted genotypes over time. We posit that intraspecific variation in *Bb* host breadth and specificity helps maintain overall species fitness in response to transmission by a generalist vector. Because pathogen genotypes vary in host breadth and result in diverse human disease manifestations, our findings indicate that a more nuanced definition of ‘host competence’ incorporating local genotype frequency is warranted.

**Significance:** Lyme disease is the most common vector-borne disease in the US with a causative agent (*Borrelia burgdorferi*) exhibiting high genetic diversity that partially correlates with human disease manifestations. Understanding the extent of host specificity in pathogens is critical for evaluating disease risk, but host specificity and mechanisms maintaining genetic diversity in Bb are unknown. We show that Bb genotypes exhibit variable host adaptation to white-footed mice and passerine birds, two common reservoir hosts, which appears to promote high intraspecific pathogen diversity. Conversely, we find limited evidence of negative frequency-dependent selection, an alternative mechanism for diversity maintenance. Our results reveal cryptic intraspecies host breadth variation and suggest that evaluating host competence depends on the frequency of host-adapted genotypes in local environments.

## Introduction

Evaluating human disease risk from zoonotic pathogens, those shared between wildlife and humans, requires characterization of pathogen traits influencing their environmental distribution and spillover potential (Sánchez *et al*. 2021). Recent meta-analyses and reviews have repeatedly identified host breadth, or the capacity to infect phylogenetically diverse species, as a key trait influencing spillover (Woolhouse & Gowtage-Sequeria 2005; Olival *et al*. 2017; Becker & Albery 2020). Greater host breadth provides pathogens more opportunities for population persistence across heterogeneous environments (e.g. variable host availability), but often invokes a fitness tradeoff as pathogens must adapt to diverse immunological selection pressures (Elena & Lenski 2003; Sexton *et al*. 2017).

Wider pathogen host breadth is intrinsically linked to increased competence of potential hosts, defined as the capacity to acquire, maintain, and transmit infection (Stewart Merrill & Johnson 2020). Though host competence is increasingly used to model human disease risk, unexplained sources of variation can limit its utility (Downs *et al*. 2019; Becker *et al*. 2020; Stewart Merrill & Johnson 2020). In particular, intraspecific genetic variation in multi-strain pathogens may influence host-pathogen interactions, preventing accurate characterization of pathogen host breadth and host competence (Balmer & Tanner 2011; Kilpatrick *et al*. 2017).

To understand the consequences of genotypic diversity on pathogen phenotypes and human disease risk, one must also characterize the evolutionary drivers that maintain diversity and the ecological context in which variation is observed. Balancing selection can maintain diverse genotypes across a bacterial population when heterogeneous immunological and physiological host environments select for different traits (Elena & Lenski 2003; Hedrick 2006; Weedall & Conway 2010). While observing evidence of balancing selection can be straightforward, identifying the eco-evolutionary processes responsible requires careful experimental design or intensive and targeted population sampling (Grenfell *et al*. 2004; Hedrick 2006; Gandon *et al*. 2008; Thrall *et al*. 2012; Koskella & Vos 2015). Thus, to evaluate diversity patterns and evolutionary drivers of pathogen traits, it is crucial to sample pathogens across endemic hosts and time scales relevant to detect natural selection (AVMA 2008; Wells & Clark 2019; Woods *et al*. 2019; Becker & Han 2021).

Vector-borne pathogens are increasingly responsible for emerging infectious diseases (Jones *et al*. 2008; Swei *et al*. 2020) and may provide unique insight into the links between the evolution of host breadth and ecology of human disease risk. *Borrelia burgdorferi* sensu stricto (hereafter *Bb*) is a spirochete bacterium that causes 476,000 human cases of Lyme disease (LD) in the United States annually (Kugeler *et al*. 2021). LD is the most common vector-borne disease in the US and continues to increase in geographic range and case numbers (Radolf *et al*. 2020; CDC 2022), particularly in the Northeast and Midwest US, where *Bb* circulates primarily in small mammal and bird species via the generalist tick vector *Ixodes scapularis* (Halsey *et al*. 2018). Despite the increasing toll of tick-borne pathogens on human health, fundamental questions about their basic biological traits, including host breadth, and the role of immune environments in structuring underlying selection mechanisms, remain unanswered (Kilpatrick *et al*. 2017; Schwartz *et al*. 2021; Tsao *et al*. 2021).

Genetic studies of *Bb* regularly identify elevated genetic diversity and signatures of balancing selection at the pathogen’s outer surface protein C (*ospC*) locus in natural populations of infected ticks, with more than 25 known alleles (Qiu *et al*. 1997; Wang *et al*. 1999; Barbour & Travinsky 2010; Brisson *et al*. 2012). The OspC protein is required for host infection (Tilly *et al*. 2006) and *ospC* alleles are often used to distinguish among strains with variable phenotypes, including a subset deemed human-invasive strains (HIS) that exhibit more severe pathology in humans (Lagal *et al*. 2006; Dykhuizen *et al*. 2008; Strle *et al*. 2011). The eco-evolutionary drivers maintaining *ospC* variation and implications for human disease risk remain a focus of active debate (O’Keeffe *et al*. 2020). Two competing hypotheses exist, centered around distinct host-pathogen interactions. The first suggests *Bb* diversity is maintained via multiple niche polymorphism (MNP) in which genotypes exhibiting host-adapted phenotypes segregate across different host species (Brisson & Dykhuizen 2004; Mechai *et al*. 2016). The second predicts that negative frequency-dependent selection (NFDS) occurs via host antibodies, which iteratively induce fitness costs on common genotypes within the local population, driving temporal fluctuations in genotype frequency (Swanson & Norris 2008; Haven *et al*. 2011).

Here we present the most comprehensive long-term study of *Bb ospC* diversity to date examined across two divergent reservoir host types, passerine birds and white-footed mice, as well as tick vectors in the United States to date. By leveraging long-read amplicon sequencing of the *ospC* locus, we reveal intraspecific variation in host adaptation phenotypes that yield strong support to the MNP hypothesis, while the absence of strain dominance shifts indicates a minor role for NFDS in maintaining strain diversity. Our results shed light on the drivers of balancing selection and the evolution of host breadth for this emerging zoonotic pathogen.

## Results

### Genotypic community structure

To understand the distribution and diversity of *ospC* major groups (oMGs, hereafter described as genotypes), we sequenced the *ospC* gene from 553 white-footed mice, 92 passerine birds (11 species, Table S1), and 628 individual nymphal *Ixodes scapularis* ticks. In total, we assigned 696,453 HiFi reads to 21 genotypes (Supplementary File 1) and sequencing depth per sample varied between 20 and 9874 reads. Coinfection with multiple genotypes was common across all samples (52.8%). Individual genotype richness ranged from 1 – 16, and was lower on average in mice (α=1.77) than in birds (α=2.59) and intermediate in nymphal ticks (α=2.30). Sequencing depth showed a correlation with genotype richness only at shallower coverage below 100 reads (p < 0.001; Figure S1). We used Hill numbers to evaluate diversity profiles (Alberdi & Gilbert 2019) and found that *Bb* diversity and evenness in mice are lower compared to those of birds and nymphs, despite similar richness across populations (Figure S2).

We identified three novel genotypes. One was designated subtype Cj and shares 97.3% sequence identity with *Bb* type C, but contains a 75bp region that is fully identical with *Bb* type J – evidence of a recent recombination event (see below). Another genotype exhibited >8% sequence dissimilarity with all known oMG genotypes and was designated type J3, which was found almost exclusively in birds (present in 0.2% of *Bb* infected mice compared to 17.4% of *Bb* infected birds). A third genotype was quite divergent with type T being its closest relative (11.2% dissimilarity), but matched *Borrelia kurtenbachii* (*B.kurt.*) *ospC* at 99.8%. *B.kurt.* is a recently described genospecies and close relative of *Bb* that infects mammals exclusively (Margos *et al*. 2010). *B.kurt.* was the only genotype never identified in birds (Table S1) and exhibited a distinct lack of co-occurrence with other genotypes in mice (Figure 3), indicating a low frequency of mixed infection between *Bb* and *B.kurt* among mice (4.8% of *Bb* infected mice).

**Figure 1.**
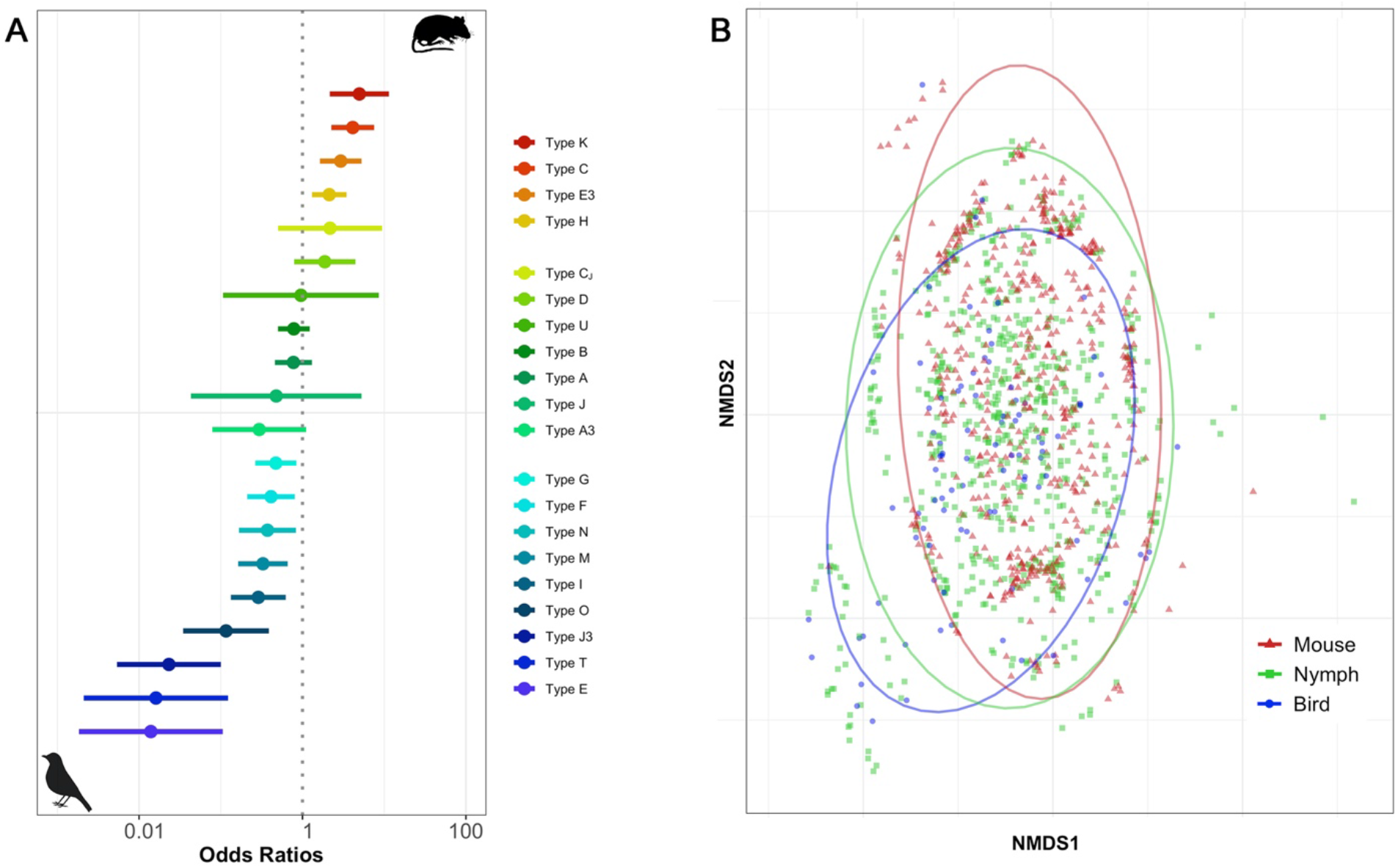
Host associations among *ospC* genotypes. (A) Log odds of infection with each genotype in mice compared to bird hosts. Significant associations exhibit confidence intervals that do not overlap the dotted line. (B) NMDS plot of individual mouse, nymph, and bird genotype communities with 95% confidence ellipses.

**Figure 2.**
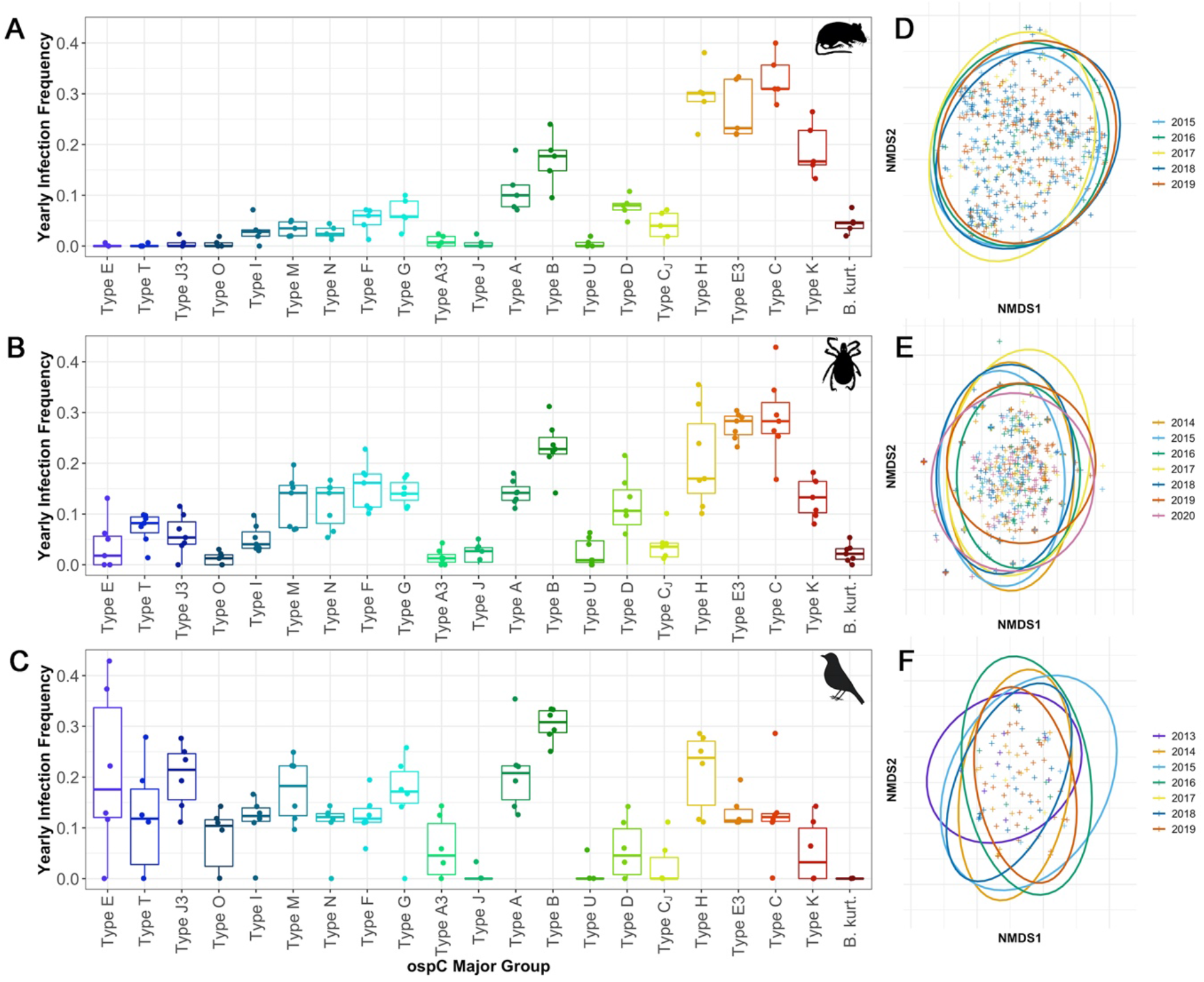
Yearly variation in genotype frequency and individual community structure. Boxplots of ospC genotype frequency distributions observed in mice (A), nymphs (B), and birds (C). NMDS plots of individual communities separated by year among mice (D), nymphs (E), and birds (F) with 95% confidence ellipses. Boxplot colors reflect strength of host-association shown in Figure 1.

**Figure 3.**
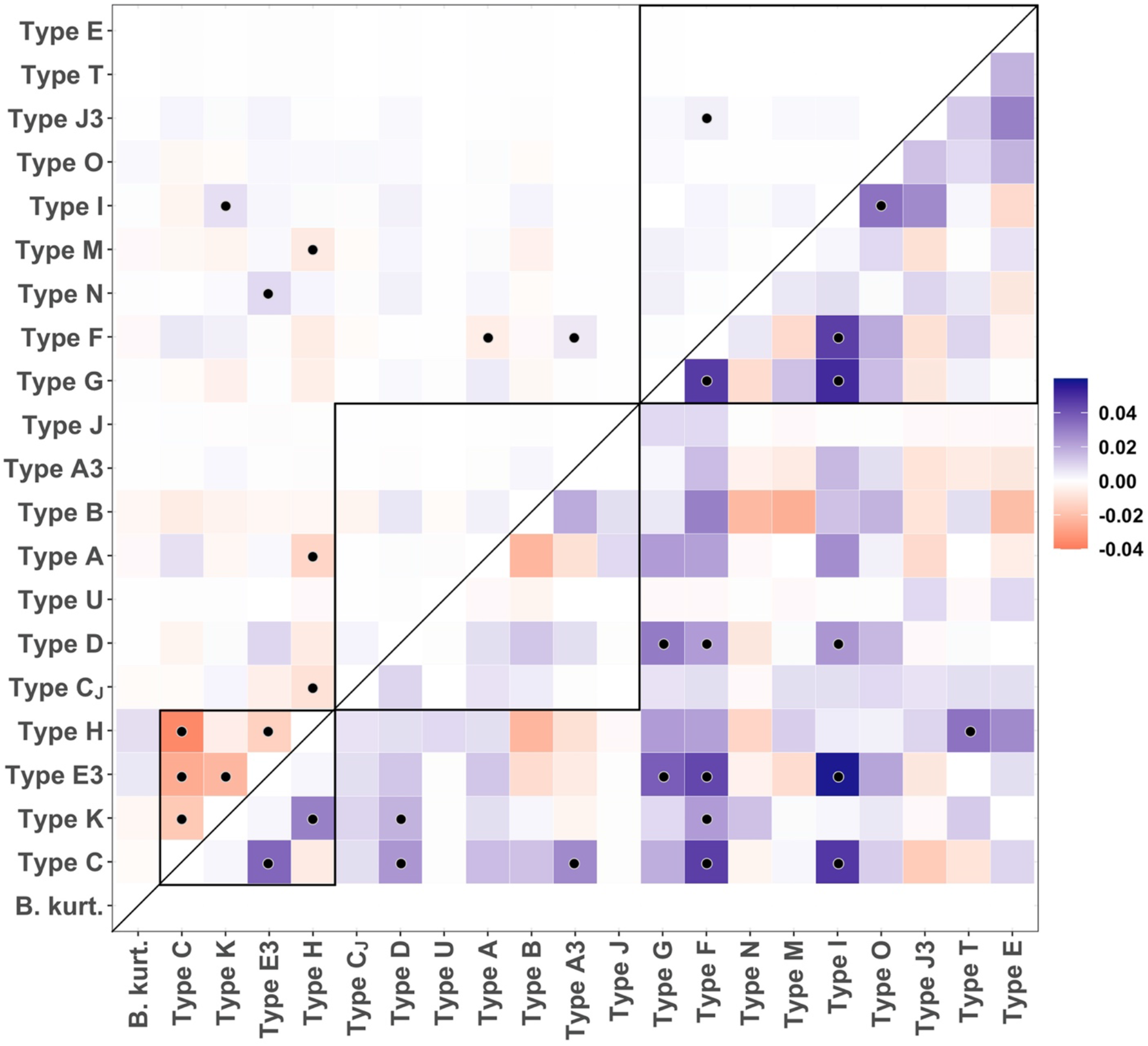
Heatmap of effect size for cooccurrence probability estimates for ospC genotypes found in individual mammal (top triangle) or bird hosts (bottom triangle). Color indicates the strength of the effect (blue = positive, red = negative), and black dots indicate significant associations (p < 0.05). Black boxes within the heatmap surround the putative mouse-adapted genotypes (bottom), generalist genotypes (middle), and bird-adapted genotypes (top).

### Evidence supporting MNP mechanisms in hosts

If genotypic diversity in *Bb* communities is maintained by MNP, we expect significant associations between genotypes and hosts, with nymphal ticks harboring all circulating genotypes. We identified strong patterns of host association among the 20 *Bb* genotypes found in this study (Figure 1A, Table 1). Four genotypes (Types C, E3, H, K) were significantly more likely to be found in rodent compared to avian hosts, whereas nine genotypes (Types E, F, G, I, M, N, O, T, CJ) followed the opposite taxonomic trend. The remaining seven genotypes (Types A, A3, B, D, J, U, J3) were not associated with either host taxon. We hereafter refer to these genotypes as mouse-adapted, bird-adapted, and generalist, respectively. The signal of host association among the most strongly bird-adapted genotype (type E), which had merely a 0.99% probability of infecting a mouse (OR = 0.01), is stronger than that of the most mouse-adapted genotype (type K), which maintains a 16% probability of infecting a bird (OR = 0.20, when mouse is set as the reference). No *Bb* genotypes were found exclusively in a single host taxon (Table S1). The dominant genotype, accounting for the largest proportion of reads, in an individual was adapted to that host taxon in the majority of observations, occurring in 376 of 553 individual mice (68.0%) and 51 of 92 individual birds (55.4%).

**Table 1.** Log odds ratio of strain-specific infection in mouse host and tick vector relative to bird hosts. Significant odds ratios are in bold and 95% confidence intervals provided in parentheses.

| *ospC*<br>Genotype | Mouse |  | Tick |  |
| --- | --- | --- | --- | --- |
|  | Odds Ratio | p-value | Odds Ratio | p-value |
| K | **4.97 (2.16 – 11.43)** | <0.001 | **2.32 (1.00 – 5.37)** | 0.050 |
| C | **4.12 (2.25 – 7.53)** | <0.001 | **2.74 (1.50 – 4.99)** | 0.001 |
| E3 | **2.94 (1.63 – 5.28)** | <0.001 | **2.31 (1.29 – 4.14)** | 0.005 |
| H | **2.13 (1.31 – 3.46)** | 0.002 | 1.10 (0.67 – 1.80) | 0.705 |
| CJ | 2.17 (0.50 – 9.44) | 0.300 | 1.98 (0.47 – 8.45) | 0.354 |
| D | 1.88 (0.79 – 4.44) | 0.153 | 2.02 (0.87 – 4.70) | 0.103 |
| U | 0.96 (0.11 – 8.60) | 0.968 | 0.16 (0.06 – 0.43) | <0.001 |
| B | 0.78 (0.50 – 1.23) | 0.285 | 0.84 (0.55 – 1.29) | 0.423 |
| A | 0.77 (0.46 – 1.31) | 0.338 | 0.70 (0.42 – 1.17) | 0.173 |
| J | 0.48 (0.04 – 5.28) | 0.546 | 2.47 (0.33 – 18.82) | 0.382 |
| G | **0.47 (0.26 – 0.84)** | 0.011 | 0.84 (0.50 – 1.41) | 0.505 |
| F | **0.41 (0.21 – 0.81)** | 0.010 | 1.12 (0.63 – 2.00) | 0.700 |
| N | **0.37 (0.17 – 0.83)** | 0.016 | 1.24 (0.63 – 2.44) | 0.529 |
| M | **0.33 (0.16 – 0.66)** | 0.002 | 0.88 (0.49 – 1.59) | 0.680 |
| A3 | **0.30 (0.08 – 1.11)** | 0.071 | 0.41 (0.13 – 1.30) | 0.130 |
| I | **0.29 (0.13 – 0.62)** | 0.002 | **0.45 (0.23 – 0.89)** | 0.021 |
| O | **0.12 (0.03 – 0.39)** | <0.001 | **0.16 (0.06 – 0.43)** | <0.001 |
| J3 | **0.02 (0.01 – 0.10)** | <0.001 | **0.32 (0.18 – 0.56)** | <0.001 |
| T | **0.02 (0.00 – 0.12)** | <0.001 | **0.16 (0.06 – 0.43)** | <0.001 |
| E | **0.01 (0.00 – 0.11)** | <0.001 | **0.21 (0.11 – 0.41)** | <0.001 |

Individual hosts or ticks are represented by their genotypic community, composed of one or more genotypes and their ranking within the individual, defined by relative sequencing depth. To understand similarity among individual genotypic communities among the populations of mice, birds, and ticks we used non-metric multidimensional scaling (NMDS). NMDS revealed moderate separation between genotypic communities typically found in each population, with birds and mice clustering further away from one another, and ticks forming an intermediate cluster with individual communities that share genotypic community characteristics with both host groups (Figure 1B). Separately, we used a pairwise similarity test of genotype overlap among populations and found that both mammal genotype and bird genotype communities were more similar to nymph genotype communities (0.904 and 0.883, respectively), than they were to each other (0.672). We observed some variation in the frequency of genotype infection across *Bb* infected bird species (Table S3) but GLMs differentiating between commonly infected bird species (Carolina Wren, Common Yellowthroat, and American Robin) indicated that most genotypes exhibited parallel responses compared to white-footed mice (Figure S3). We thus combined bird-derived data in our analyses.

### Limited evidence of NFDS mechanisms in hosts

If genotypic diversity in *Bb* is maintained by NFDS, we would expect significant variation in the dominant genotype and shifting genotypic frequencies within host populations over time. We first examined genotype frequency and similarity of individual genotype communities within populations across years (Figure 2). For mice, where sample sizes among years were consistently high, the distributions of annual genotype frequencies exhibited low dispersion over time (Figure 2A, Figures S4). Similar patterns were observed among nymphal ticks and bird hosts, though smaller sample sizes and variation among bird species led to more variation in frequency distributions (Figure 2A, Figure S5, S6).

We used analysis of similarities (ANOSIM) to test for significant differences in the composition of individual communities within and between years for each taxonomic population, separating analyses across sampling sites for mice and nymphal ticks. We found significant differences in mouse genotype communities across years at only one of the three sampling sites (RH: p = 0.047, NR: p = 0.09, EI: p =0.673), but not among bird infection communities, which were sampled at multiple sites across the island (p = 0.85). Pairwise post-hoc comparisons revealed significant differences in mice at RH only between 2015 and 2016, 2018, and 2019. In contrast, questing nymphs exhibited significant yearly differences in their infection community across all three sites (RH: p = 0.001, NR: p = 0.001, EI: p =0.004). Graphic examination using NMDS plots revealed strong and consistent overlap for within-year variation in genotype communities across populations of mice, birds, and ticks (Figure 2B).

### Divergent genotype co-occurrence patterns among hosts

Competitive or facilitative interactions among genotypes within hosts may influence their probability of co-occurrence within the same individual host community. For each population of mice, birds, and ticks we compared the observed pairwise co-occurrence of genotypes within individual infection communities to the random distributions predicted by the total number of genotype occurrences in that respective population. We found contrasting patterns in the number and directionality of significant genotype pairs across mice and birds (Figure 3). Among mice, we observed 13 significant pairwise correlations, including multiple negative correlations among the four mouse-adapted genotypes (i.e. pairs of genotypes were observed together less frequently than expected). Among birds, we observed a greater total number of significant correlations (n =19) than in mice, all of which were positive (i.e. pairs of genotypes were observed together more frequently than expected), with no consistent relationship among bird-adapted genotypes. Among nymphs, we observed both positive and negative significant correlations among genotypes, with strong negative interactions between those genotypes with the strongest signals of adaptation to birds and mice, suggesting ticks rarely acquire both types of genotypes through a larval bloodmeal (Figure S7).

We examined whether genetic dissimilarity or phylogenetic network distance among genotype pairs predict the observed co-occurrence effect size using linear models. No significant relationship was found between either predictor variable and co-occurrence in mice (p = 0.097 and p = 0.096) or nymphs (p =0.087 and p = 0.082), but a significant negative relationship between both predictors and co-occurrence was found in birds (p = 0.007 and p =0.008, Figure S8), indicating that co-occurrence was less frequent when genotype pairs were more genetically divergent.

### Individual mouse genotype community dynamics

The genotype community of individual hosts may change over time due to the host’s ability to clear infections by specific genotypes and due to sequential introductions through multiple nymphal tick bites. To understand how patterns of genotype host adaptation influence individual-level infection dynamics, we used a multistate Markov model to dissect mice genotype infection in mice sampled multiple times within a single year. We characterized the state of infection based on the identity of the dominant genotype (i.e. greatest sequencing depth) at each sampling time, specifying three possible states: 1) uninfected, 2) infected with a mouse-adapted genotype, 3) infected with a non-mouse-adapted genotype (i.e. generalist or bird-adapted). We found that mice in any initial state are more likely to transition to a mouse-adapted than a non-mouse-adapted genotype infection (Table 2). We also found that the mean persistence (i.e. mean sojourn time) of infections was nearly three times greater in mouse-adapted than that in non-mouse-adapted genotypes in mice (27.6 days vs. 9.5 days; Table S4). Yet, when the state of an uninfected mouse changes, we found it was twice as likely to become infected with a non-mouse-adapted genotype than a mouse-adapted genotype (Table S5). The dominant genotype was significantly more likely to change with increasing time since initial capture (Figure S9) but was not influenced by individual characteristics or tick burden (Table S6). Together, our model suggests that while infections with non-mouse adapted genotypes are common in mice, these genotypes exhibit weak persistence and are often replaced with infections dominated by mouse-adapted genotypes.

**Table 2.** Transition probabilities between infection states in white-footed mice estimated over a 4-week period with the multistate Markov model. Columns and rows represent starting and transition states, respectively. 95% confidence intervals are provided in parentheses.

|  | **Uninfected** | **Adapted** | **Non-Adapted** |
| --- | --- | --- | --- |
| Uninfected | 0.722 (0.664 - 0.765) | 0.346 (0.277 - 0.258) | 0.167 (0.121 - 0.237) |
| Adapted | 0.164 (0.131 - 0.212) | 0.501 (0.395 - 0.580) | 0.336 (0.252 - 0.42.7) |
| Non-Adapted | 0.114 (0.088 - 0.146) | 0.152 (0.111 - 0.205) | 0.497 (0.401 - 0.592) |

### Evolutionary patterns among genotypes

We examined evolutionary patterns at the *ospC* locus. The Neighbor Net phylogenetic network revealed long branches separating each *ospC* type, with pronounced reticulation at the center of the network (Figure 4A). The single exception was between type C and type J3, which are closely related (97.3% nucleotide similarity). Examination of genotype representative amino acid sequences revealed a K196Q mutation, located at the C-terminus of OspC’s 5^th^ α-helix. All mouse-adapted genotypes and all but one of the generalist genotypes exhibit a lysine at this residue, while all but one bird-adapted genotypes exhibit a glutamine (Figure 4B).

**Figure 4.**
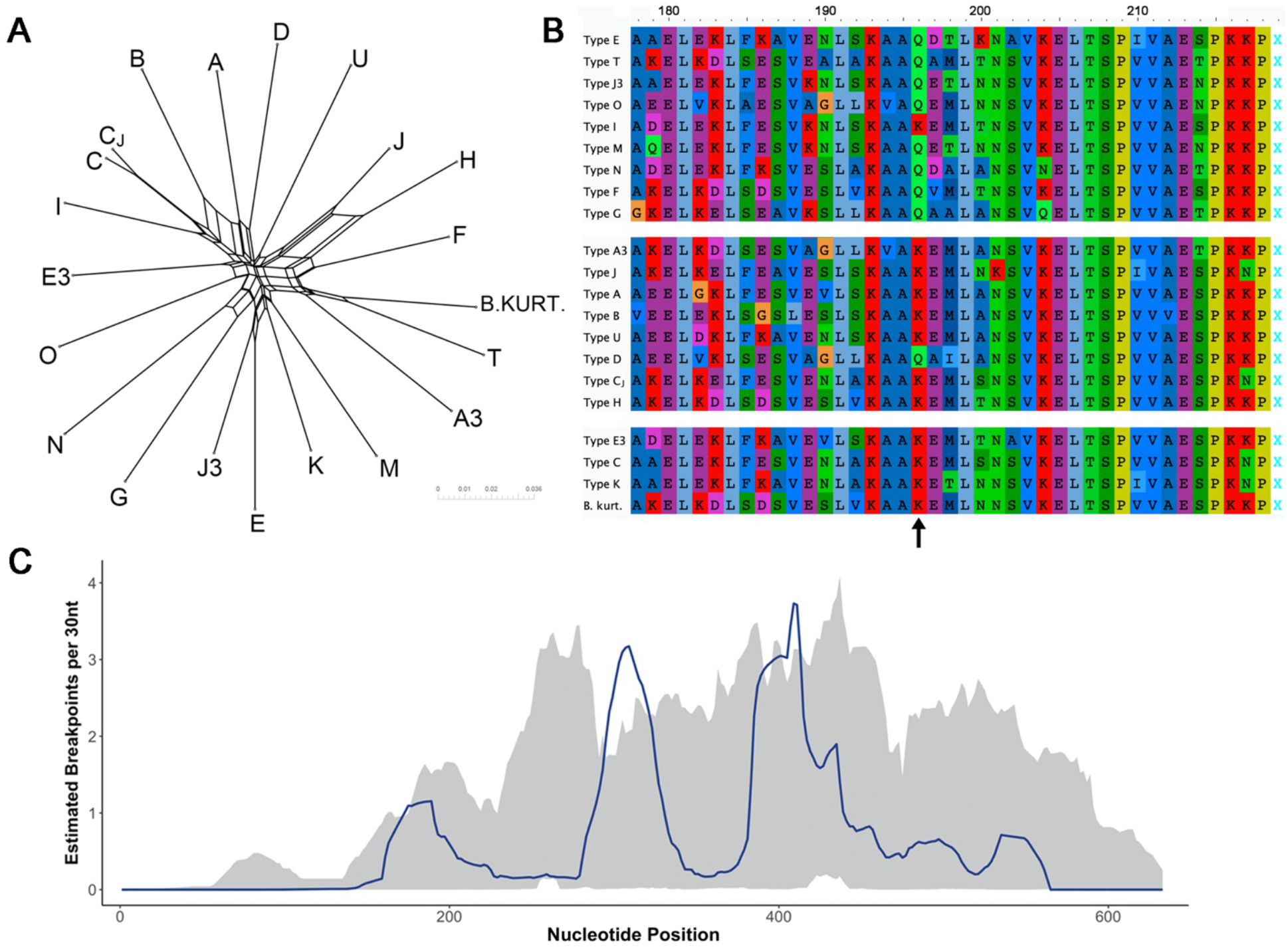
Evolutionary patterns at the *ospC* locus. (A) Phylogenetic network of 21 observed genotypes. (B) Amino acid alignment of 5’ end of *ospC* gene with arrow identifying the K196Q mutation. Genotypes are separated by phenotypic groups: bird-adapted (top), generalist (middle), mouse-adapted (bottom). (C) Recombination breakpoint estimation (blue line) across *ospC*. Shaded area represents 95% confidence intervals for local hot/coldspot test. Areas where observed counts exceed confidence interval are inferred recombination hotspots.

Using a suite of recombination detection tools, we identified 11 intralocus recombination events across the 21 *ospC* types, each of which was detected by at least three independent methods (Table S7). Analysis of breakpoints suggested two major recombination hotspots around nucleotide positions 300 and 400, with lower confidence hotspots in positions 180 and the 500-550 region (Figure 4C).

## Discussion

A critical step in evaluating human disease risk from emerging zoonotic pathogens is understanding how interactions with endemic reservoir hosts shape a pathogen’s genetic and functional diversity, in particular its host breadth and ability to infect new species (Karesh *et al*. 2012; Olival *et al*. 2017). Here we demonstrate that *Borrelia burgdorferi* sensu stricto exhibits a spectrum of host-adapted phenotypes associated with the *ospC* alleles. Specifically, we identified several genotypes that show evidence of adaption to white-footed mice (Types C, E3, H, K) and passerine birds (Types E, F, G, I, M, N, O, T, Cj), while a group of genotypes (Types A, A3, B, D, J, U, J3) exhibit no significant association with either host (i.e. are generalists). Importantly, the prevalence of *Bb* genotypes in ticks, including human invasive strains (HIS), closely reflects genotypic frequency in the local host community. These results reveal cryptic intraspecific variation in pathogen host breadth, suggesting that host competence for *Bb*, as well as human disease risk and Lyme Disease severity, is dependent on the relative prevalence of locally circulating genotypes.

The evolutionary drivers maintaining *ospC* polymorphisms in nature have been debated in recent decades. Some field-based studies have suggested MNP drives *Bb* diversity through evidence of genotype-host associations defined by genotypes (Brisson & Dykhuizen 2004; Vuong *et al*. 2014) or multi-locus sequence typing (Mechai *et al*. 2016), though others found no such pattern (Hanincová *et al*. 2006). Lab-based studies have observed variable infection establishment, persistence, and transmission of *Bb* genotypes within and among host species (Wang *et al*. 2001, 2002; Derdáková *et al*. 2004; Hanincová *et al*. 2008; Lin *et al*. 2022). Yet, other short-term field studies (i.e. up to 3 years) observe shifting frequencies of *Bb* genotypes in tick vectors consistent with balancing selection mediated by NFDS (Qiu *et al*. 1997; Wang *et al*. 1999; States *et al*. 2014). Pervasive local recombination, revealed by recent *Bb* comparative genomics, appears to drive nucleotide and antigenic diversity at *ospC*, and NFDS is proposed as a parsimonious explanation for the maintenance of this variation, particularly if recombination disrupts host-pathogen allele-level co-adaptation (Haven *et al*. 2011; Qiu & Martin 2014; Schwartz *et al*. 2021). Our results, spanning multiple host taxonomic groups and nymphal ticks over up to 7 years, provide strong evidence supporting MNP in structuring *Bb* genetic diversity. While we observed little evidence of temporal shifts in population-level genotype frequency, as predicted by NFDS, it is possible that antigenic exclusion in individual hosts accounts for minor variation among host-adapted genotypes.

While MNP appears to partially maintain *Bb* diversity through genotype-specific host adaptation, variable OspC proteins are also clearly under selection for distinct antigenic epitopes that limit cross-reactivity (Baum *et al*. 2013). Since processes governing initial establishment of infection through occur before the production of antibodies, additional selective forces must maintain diverse antigenic epitopes (Kurtenbach *et al*. 2006). We propose that limited antibody cross-reactivity serves to allow MNP processes by enabling serial susceptibility of hosts. Because hosts produce antibodies targeting specific genotypic epitopes (Probert *et al*. 1997; Baum *et al*. 2012), subsequent infections by different genotypes encounter essentially naïve hosts. Thus, short-lived infections by non-adapted genotypes should not restrict the future success of more host-adapted genotypes, improving the likelihood of transmission despite variable infection success. In the Northeast US, nymphal ticks are abundant for 6-8 weeks, several weeks before the emergence of larvae, allowing hosts to experience multiple *Bb* introductions from independent tick bites. Longer duration phenotypes then have a higher likelihood of infecting the next tick cohort’s larvae (Ogden *et al*. 2007; Haven *et al*. 2012). Indeed, variable tick phenology has been shown to be associated with infection persistence (Gatewood *et al*. 2009). In our study, not only did mouse-adapted genotypes exhibit much longer persistence times in mice sampled multiple times than non-mouse-adapted genotypes, but mouse infection communities preferentially shifted mouse-adapted genotype dominance over time. While temporal shifts in *Bb* infections have previously been observed in mice and humans (Anderson & Norris 2006; Nadelman *et al*. 2012; Khatchikian *et al*. 2014), our results highlight how the combination of limited antibody cross-reactivity and variable host-adapted pathogen genotypes may provide a fitness advantage for *Bb* when hosts experience multiple independent infections.

A second mechanism maintaining genotype diversity is the lack of complete host specialization; even *Bb* genotypes with the strongest evidence for host adaptation were observed in their non-adapted hosts. This pattern may be an intrinsic aspect of the genotype phenotype, may be due to intraspecific variation in host competence, or result from facilitative interactions among genotypes that allow non-adapted genotypes to persist in the presence of host-adapted genotypes (Andersson *et al*. 2013; Stewart Merrill & Johnson 2020). Experimental studies of bacterial evolution find that increasing environmental complexity (e.g. host diversity) results in overlapping niches and imperfect specialization (Barrett *et al*. 2005). A recent model examining the interaction between MNP and NFDS in *Bb* also suggested that more diverse suites of differentially host-adapted phenotypes can coexist in a population when antigenic variation, a form of environmental heterogeneity, is high (Adams *et al*. 2021). These findings are suggestive of a role for host adaptation-related pathogen traits in maintaining pathogen fitness and persistence in infected hosts. The high genotype diversity observed in *Bb* appears to fit a ‘mass effect’ metacommunity model (Leibold *et al*. 2004; Alexander *et al*. 2012) where host-adapted genotypes are common in their respective hosts and strong propagule pressure (from infectious ticks) results in low frequency of non-adapted genotypes co-occurring with adapted ones.

### Host community composition drives patterns of Bb diversity in ticks and human disease risk

The frequency of specific *Bb* genotypes in nymphal ticks, which are the main source of human infections, matches the genotype prevalence observed in the local host community (Figure 2A). This trend is particularly important for understanding the local prevalence of human invasive strains (HIS), a subset of *Bb* genotypes including ospC types A, B, I K, and N that produce more severe human disease outcomes (Seinost *et al*. 1999; Dykhuizen *et al*. 2008; Wormser *et al*. 2008). Human disease risk appears to reflect both the presence of reservoir hosts as well as the host breadth and specificity phenotypes exhibited by local *Bb* genotypes. Further, suggests a more nuanced view of a host’s pathogen competence and their role as amplification or dilution hosts, where the infectious phenotype of the host-adapted genotype should be considered in addition to the pathogen’s prevalence in the host.

In our study, birds were the main source of ospC types I and N and harbor types A and B in high frequency, although not as high as mice. Though *Bb* infection prevalence varies among bird species (Ginsberg *et al*. 2006; Becker & Han 2021), previous work estimated that 27% of larval ticks feed on birds at our study location (Huang *et al*. 2019). Higher genotype richness and evenness were observed in bird hosts compared to white-footed mice. Further, we found that individual bird infection communities were characterized by greater genotype co-occurrence, indicating facilitation, in contrast with mammal communities that appear to exhibit only competitive interactions among genotypes. These findings may be partially explained by birds’ longer lifespan, different immunity mechanisms, larger home ranges, or other life-history traits resulting in more opportunities for infection (Rataud *et al*. 2021). Thus, our study emphasizes the role of passerine birds in maintaining *Bb* transmission and increasing human disease risk locally, in addition to their known roles as reservoir hosts and tick dispersers (Ginsberg *et al*. 2006; Brinkerhoff *et al*. 2010, 2011; Vuong *et al*. 2014; Loss *et al*. 2016; Becker & Han 2021).

### Signatures of adaptation in ospC

*Bb* genomes are characterized by the presence of paralogous lipoprotein gene families, whose members provide multiple and redundant roles throughout different stages of infection (Wywial *et al*. 2009; Caine & Coburn 2016), though o*spC* has no close paralogs (Mongodin *et al*. 2013). The OspC protein has known roles binding to multiple host ligands to promote evasion of host complement, binding to tick salivary proteins to prevent host antibody attack, and promoting host tissue colonization and dissemination (Ramamoorthi *et al*. 2005; Lagal *et al*. 2006; Bonnot *et al*. 2010; Caine *et al*. 2017; Lin *et al*. 2020). The *ospC* K196Q mutation we identified may contribute to host adaptation *Bb* phenotypes, as it was observed in 8 out of 9 bird-associated genotypes, but only in 1 out of 7 generalist genotypes, and was absent in mouse-adapted genotypes. This mutation localizes in the N-terminus of OspC’s 5^th^ α-helix, a region predicted to be important for genotype-specific antibody cross reactivity (Baum *et al*. 2013). Though *ospC* variation clearly impacts antigenic recognition, the elevated role of MNP identified here suggests it may also influence host specific adaptation. If this gene is involved in host specific complement evasion or tissue dissemination in addition to influencing antibody cross-reactivity, it would suggest that competing or complementary evolutionary pressures govern standing variation. Importantly, we posit that genotype identity alone does not dictates host adaptation phenotypes, as other loci in linkage disequilibrium with *ospC* are likely involved in conferring host-adapted phenotypes via multiple interrelated mechanisms (Dykhuizen & Dean 2004). Our results suggest genotype-host associations may be partially due to variation at *ospC* directly impacting fitness across host immune environments and should be a target of further investigation.

Studies of *ospC* variation consistently report <2% and >8% nucleotide variation within and among all oMG genotypes, respectively (Wang *et al*. 1999; Lin *et al*. 2002; Brisson & Dykhuizen 2004). This bimodal distribution is expected to reflect antigenic interactions, whereby different genotypes display distinct epitopes that limit cross-reactivity of host antibodies (Probert *et al*. 1997; Earnhart *et al*. 2005). Interestingly, the genotype identified here as type C_J_ does not conform to this widespread trend, exhibiting 2.7% nucleotide difference to type C. Our recombination analysis indicated that genotype type C_J_ is the product of incorporation of the type J entire 4^th^ α-helix (residues 158-183) into type C. A similar variant (type C_KR10_) was previously reported circulating in tick populations in both the Northeast and Midwest US (Brisson *et al*. 2010). In our study, type C_J_ exhibits decreased association with white-footed mice compared to genotype type C. Together, these patterns suggest that selective pressures otherwise maintaining coinfection with divergent genotypes is relaxed for this genotype, though the mechanism involved and the potential role of differential host adaptation require further study.

## Methods

### Study site and sample collection

Nymphal ticks were collected on Block Island, RI, USA, between May and August, 2014 - 2020. Small mammals, mainly white-footed mice, were trapped in the same months from 2014 - 2019, and birds were sampled in the same months from 2013 - 2019. Grids were established for tick and small mammal sampling at three sites across the island at the Block Island National Wildlife Refuge (NR), a private property on the eastern part of the island (EI), and at Rodman’s Hollow (RH). Passerine birds were mist netted at two sites at the Ocean View Pavilion (OVP) and Bayrose cabin (BR) under the supervision of Kim Gaffett (U.S. Department of Interior Banding Permit #09636). Nymphal ticks were collected from each grid by a standard 1m^2^ drag cloth method (Falco & Fish 1992) stopping every 10m to remove attached ticks which were immediately stored in 70% ethanol until DNA extraction. Small mammals were trapped biweekly at each site for seven sessions consisting of three consecutive trap nights each, using Sherman live traps in accordance with approved Columbia University IACUC protocol (AC-AAAS6470) and scientific collections permits from Rhode Island Department of Environmental Management and town of New Shoreham, RI. Block Island supports a low diversity small mammal community (Comings 2006) dominated by *Peromyscus leucopus* (white-footed mouse), with <1% other hosts (Huang *et al*. 2019). Upon capture morphological characteristics, such as body, tail, and foot measurements, weight, sex, age, and reproduction status were obtained. Each animal was carefully examined for ticks and other ectoparasites and marked with a uniquely numbered ear tag. Ear biopsy tissue was collected and stored in 100% ethanol until DNA extraction and pathogen analysis. Engorged *I. scapularis* larvae were removed from all bird species and stored in 70% ethanol until DNA extraction and subsequent pathogen analysis.

### DNA extraction, amplification, and sequencing

Genomic DNA was extracted using the DNeasy Blood & Tissue kit optimized for QIAcube HT automation (Qiagen). Questing nymphal and engorged larval ticks from birds were dried, frozen with liquid nitrogen, and crushed with sterile pestles before extraction. Mouse ear tissue was processed using the manufacturer’s optimized protocol. Individual samples were then screened using a duplex quantitative PCR for *Borrelia burgdorferi* and *Borrelia miyamotoi* in duplicate on an ABI 7500 Fast Real-Time PCR System using TaqMan Fast Advanced Master Mix (Thermo Fisher Scientific).

For *Bb* positive samples, we used a standard PCR to amplify a region centered around the *ospC* locus. Amplification was first attempted for a 1500bp region with the following primers GGGATCCAAAATCTAATACAA (forward) and CCCTTAACATACAATATCTCTTC (reverse). If samples failed to amplify due to DNA degradation, we attempted amplification of a smaller 750bp region that contained the entire *ospC* locus using the following primers GAGGCACAAATTAATGAAAAAGAA (forward) and GACTTTATTTTTCCAGTTACTTTTT (reverse). Both primer sets were designed to target conserved sequences identified across 18 *Bb* genotypes downloaded from GenBank (accession codes provided in Table S8). The PCR protocol included 30 s of denaturation at 98°C followed by 35 cycles of denaturation at 98°C for 10 s, annealing at 60°C for 10 s, and extension at 72°C for 1 min for the longer fragment or 30 s for the shorter fragment, followed by a final extension for 5 min at 72°C.

Each sample was amplified using a unique set of barcoded forward and reverse primers to enable pooling and downstream sample demultiplexing (Tables S9, S10). Amplification success was assessed by running 2 µl PCR product aliquots on a 1.5% agarose gel. Successful PCR products were cleaned with the ProNex Size-Selective Purification System (Promega) then quantified with a Qubit 3.0 Fluorometer using the dsDNA High Sensitivity Assay kit (Thermo Fisher Scientific) or Fragment Analyzer using the HS NGS Fragment kit (Agilent). Products were pooled at equimolar concentrations and libraries were constructed for sequencing using the single-molecule real-time (SMRTbell) sequencing Express Template Prep Kit 2.0 with an extended DNA damage repair step, then sequenced with the Pacific Biosciences Sequel I platform at the Icahn School of Medicine at Mt. Sinai Genomics Core Facility (New York, NY).

### Sequence clustering and identification of genotypes

High fidelity (HiFi) reads were generated from subreads of each zero-mode waveguide (ZMW) using the circular consensus sequencing (*ccs*) tool (https://github.com/PacificBiosciences/ccs) and demultiplexed using *Lima* v.2.2.0 (https://github.com/PacificBiosciences/barcoding) with the following parameters: --ccs --dump-removed --dump-clips --guess 80 --min-score 80 --split --output-handles 600. To identify the distribution of *Bb* genotypes among samples we first created a FASTA containing all demultiplexed reads and used *PROKKA* gene annotation software (Seemann 2014) to isolate full-length gene sequences. We used *vsearch* v2.18.0 toolkit (Rognes *et al*. 2016), a free and open-source version of the popular *usearch* OTU clustering software, to cluster all sequenced ospC genes and 32 reference sequences with known ospC types (Table S11), using --id 0.98. Of 21 read clusters with a minimum of 200 reads (all other clusters contained <5 reads), 18 clustered with known *ospC* types. We subjected the centroid sequences representing any genotypes without known matches to BLASTN against the full Genbank nucleotide database for further identification. Any novel genotype with >8% nucleotide dissimilarity to known genotypes was named according to the sequential list of all known genotypes (i.e. starting with J3), while those with <8% nucleotide dissimilarity were given a subtype designation according to its closest match (i.e. type C_J_). A novel genotype most closely matching the *ospC* of *Borrelia kurtenbachii* was named simply “B.kurt.” but not given a designated type name. Finally, we used SRST2 (Inouye *et al*. 2014) to assign reads from each demultiplexed sample represented by a minimum of 20 HiFi reads to one of the 21 identified *ospC* types using the following parameters: -- min_depth 1 --prob_err 0.005 --max_divergence 2.5.

### Statistical analyses

Statistical analyses and data filtering were conducted in R v4.1.1 unless otherwise noted (R Core Team 2021). For each sampled host or nymphal tick, we first filtered any genotype represented by less than three individual HiFi reads. We then rank transformed genotype abundances for each sample, as PCR-based amplification preferentially amplifies the most common substrate, skewing relative differences between amplified products. We evaluated diversity profiles across mammal, bird, and nymph genotype communities using Hill numbers and further characterized the extent of genotype overlap between genotype communities with the Sorensen index, both implemented with the SpadeR R package (Chao *et al*. 2016). We also examined correlations between sequencing depth on genotype richness across samples using linear models.

Because the *ospC* locus is heavily influenced by recombination, individual phylogenetic trees do not accurately represent the evolutionary relationships among genotypes. Thus, we built a phylogenetic network using Neighbor-Net (Bryant & Moulton 2004) after first aligning representative full length *ospC* sequences from each genotype (i.e. centroids of USEARCH clusters) as translated amino acids with *MAFFT* (Katoh & Standley 2013).

We investigated patterns of co-occurrence between all pairwise genotype combinations across individual mice, birds, and ticks, separately using the *cooccur* R package, which evaluates the predicted and observed probability of genotype co-occurrence within individuals given their abundances in the dataset (Griffith *et al*. 2016). To better interpret patterns of co-occurrence among genotypes we built linear models in which either nucleotide dissimilarity, as calculated by Clustal-Omega (Sievers & Higgins 2014), or phylogenetic network distance, was used to predict the effect size for each pair of genotypes.

We tested for evidence of recombination among genotypes using the RDP5 analysis suite across 30bp sliding windows using the following methods: RDP, GENECONV, BOOTSCAN, MAXCHI, CHIMERA, SISCAN. Putative breakpoints were accepted if detected across three or more methods independently (Martin *et al*. 2021).

To test for evidence of host-adapted associations between specific genotypes and mammalian or avian hosts we used binomial (logarithmic) generalized linear models (GLMs), specifying infection with an individual genotype as the response and sample type (i.e. mammal, bird, or nymph) as the predictor, setting “bird” as the reference to highlight differences between birds and mice. To account for detection bias introduced at low sequencing depth we included an offset term for samples with <100 reads (otherwise set to 1; Figure S1). A genotype was considered host-adapted if it displayed a significant association (α = 0.05) with mammalian or avian hosts. We visualized differences in genotype communities among hosts and ticks using nonmetric multidimensional scaling (NMDS) with the *metaMDS* function from the *vegan* R package (Oksanen *et al*. 2020). To examine differences among bird species, we conducted binomial GLMs as described above, but restricted bird samples to species with sample size greater than 10 (Carolina wren: *n* = 28, Common yellowthroat: *n* = 20, American robin: *n* = 16, all others ≤ 8) and included species as the predictor variable, setting “mouse” as the reference.

To test for evidence of temporal variation in genotype frequency, as expected under NFDS, we first calculated the frequency of each genotype within each sample type for each year, then used boxplots to visualize the distributions. We also plotted changes in frequency over time for each genotype at each site. To assess changes in community structure we used Analysis of Similarities (ANOSIM) tests from the *vegan* R package (Oksanen *et al*. 2020). For each sample type, communities across each year were compared using Bray-Curtis dissimilarity matrices and run with 999 permutations. If this global test was significant, we tested pairwise yearly combinations for post-hoc identification of significant pairs. We visualized differences in genotype communities by year using NMDS as described above.

To evaluate the dynamics of genotype community transitions and persistence at the individual host scale we examined mice that were sampled multiple times within a single year (n = 383) using a multi-state Markov (MSM) model, implemented with the *msm* R package (Jackson 2011). We established three infection states reflecting whether the mouse exhibited no *Bb* infection, *Bb* infection dominated (i.e. read depth rank = 1) by a mouse-associated genotype (type C, E3, H, K), or *Bb* infection dominated by a non-mouse-associated genotype. We estimated the transition rates between state pairs by maximum likelihood using ‘nlm’ optimization. We then extracted transition probabilities over a 4-week period, the mean sojourn time (i.e. persistence) for each state, and the probability of the next state given each starting state. To understand what factors influence genotype community changes we also used a binomial generalized linear mixed model (GLMM) to model the probability of a change in identity for a mouse’s dominant genotype as a function of time (days since first sampling), genotype richness, sex, maturity, and tick burden, using the site and year of capture as random variables.

## Acknowledgements

The authors thank all the Block Island field crews who collected samples between 2013 - 2020 and Ella Steiger for laboratory work. We thank Kim Gaffett and the Nature Conservancy staff on Block Island for their continued support. We also thank Mathilde Cordellier, Karina Garcia, and Nina Skinner for their contributed silhouettes to phylopic.org. This study was supported by the National Institute of General Medical Sciences, National Institutes of Health, Ecology and Evolution of Infectious Disease Program (R01 GM105246; United States), and National Science Foundation (IOS 174995, 1755286, 1755370; United States).

---

**Figure S1.**
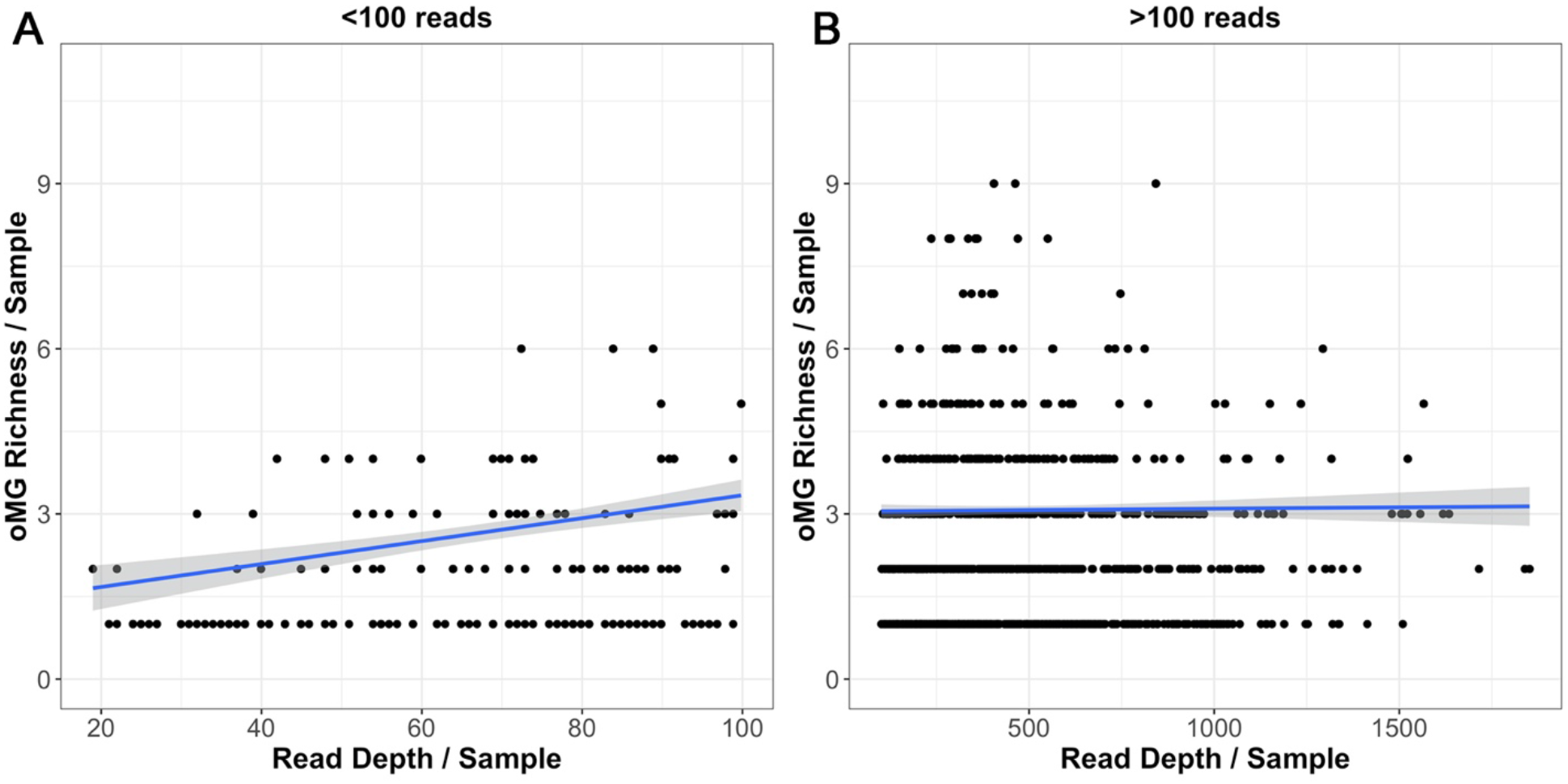
Sequencing depth bias observed for samples represented by fewer than 100 reads (A) but absent for samples represented by more than 100 reads (B).

**Figure S2.**
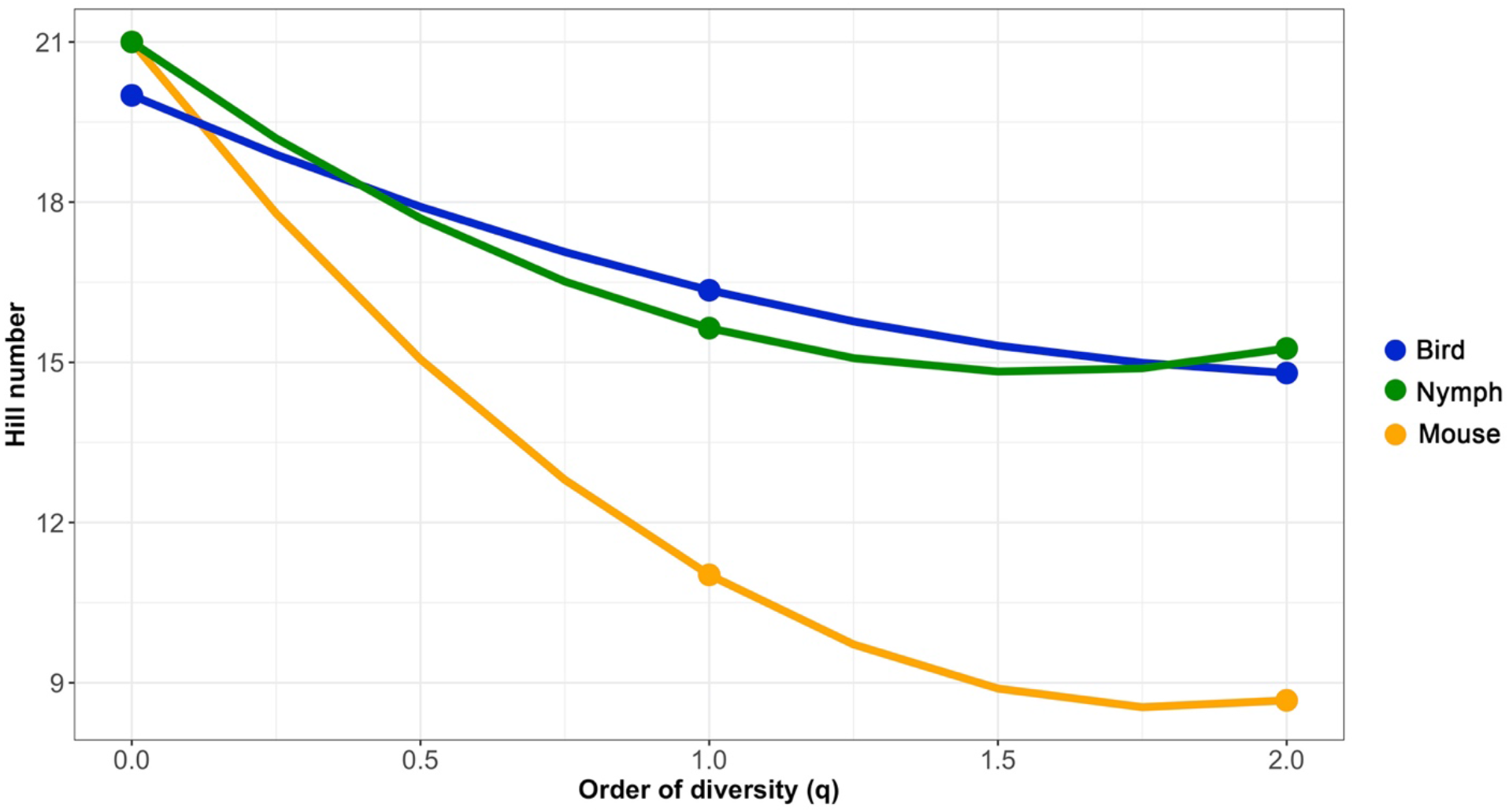
Diversity profiles using Hill numbers for genotype communities observed within mice (orange), birds (blue), or nymphs (green). Flatter profiles represent greater evenness and steeper slopes represent increasing unevenness. q = 0 is equivalent to community richness, while a q = 1 is equivalent to the Shannon index of diversity, and a q = 2 is equivalent to the multiplicative inverse of the Simpson diversity index.

**Figure S3.**
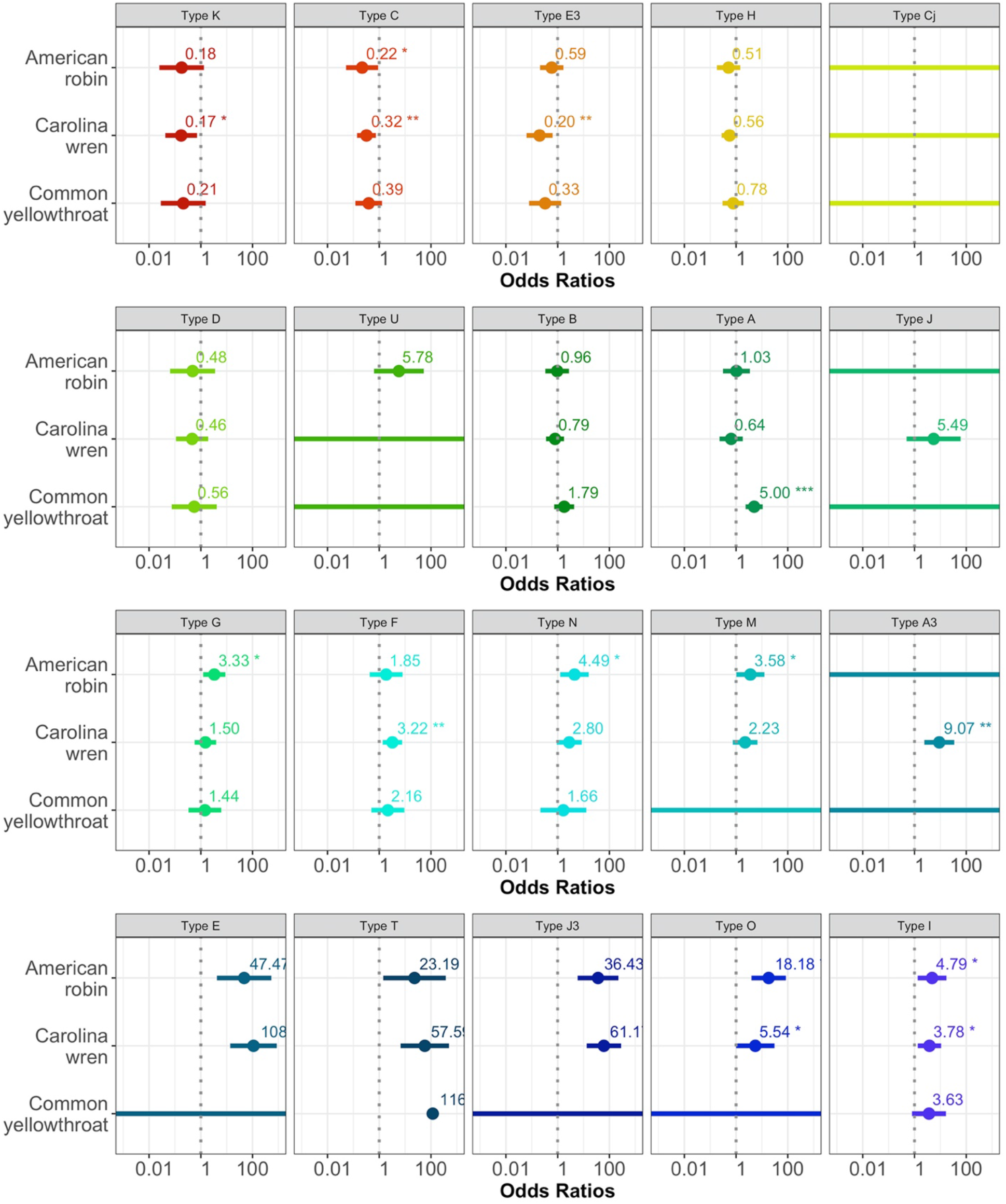
Bird species specific binomial GLMs for each *Bb* genotype. Odds ratios are relative to genotype infection in mice. Colors represent strength of host association observed when bird species were treated as a single taxa group, as labeled in Figure 1.

**Figure S4.**
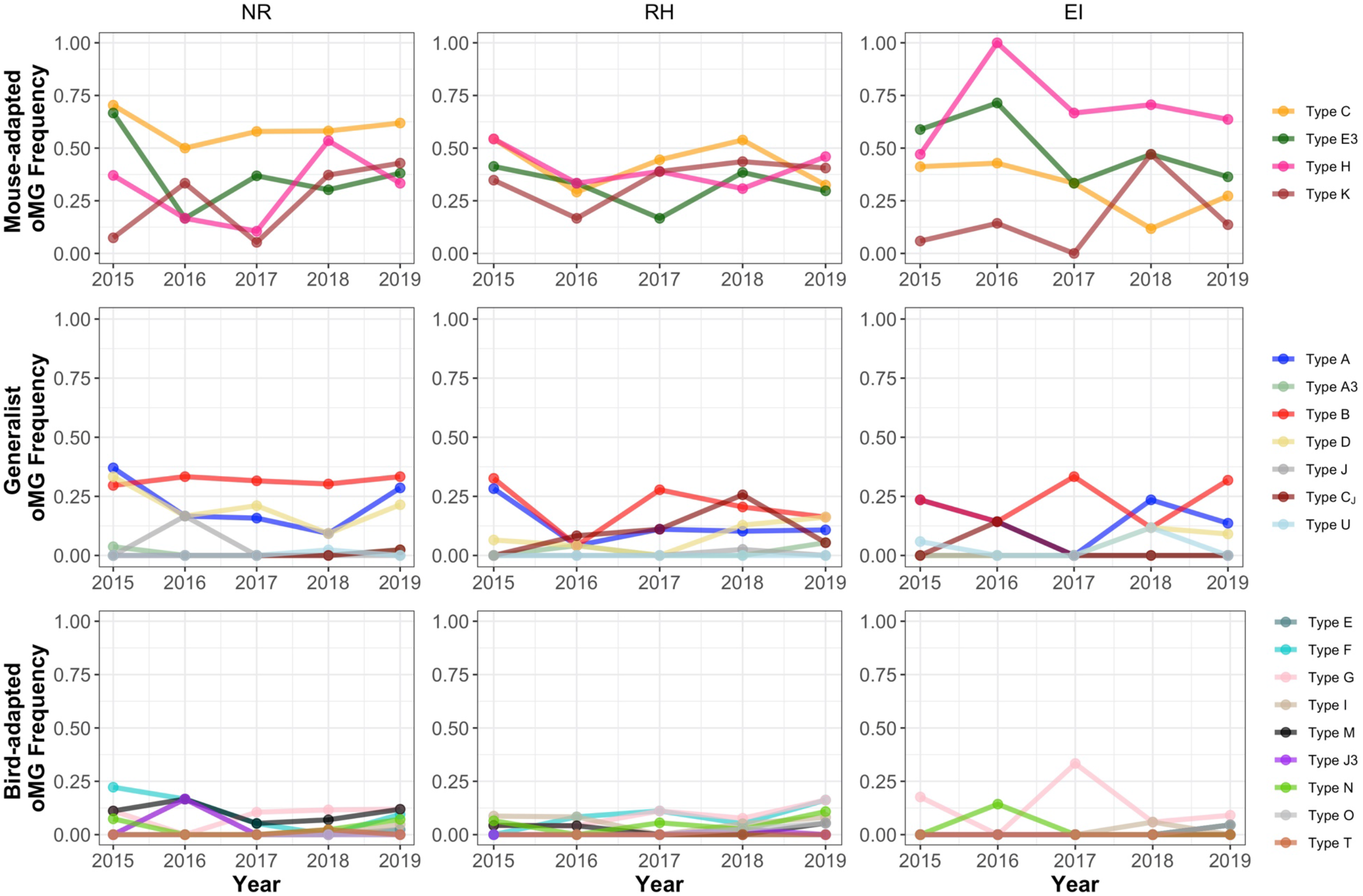
Yearly frequency of individual genotypes in mice across sites.

**Figure S5.**
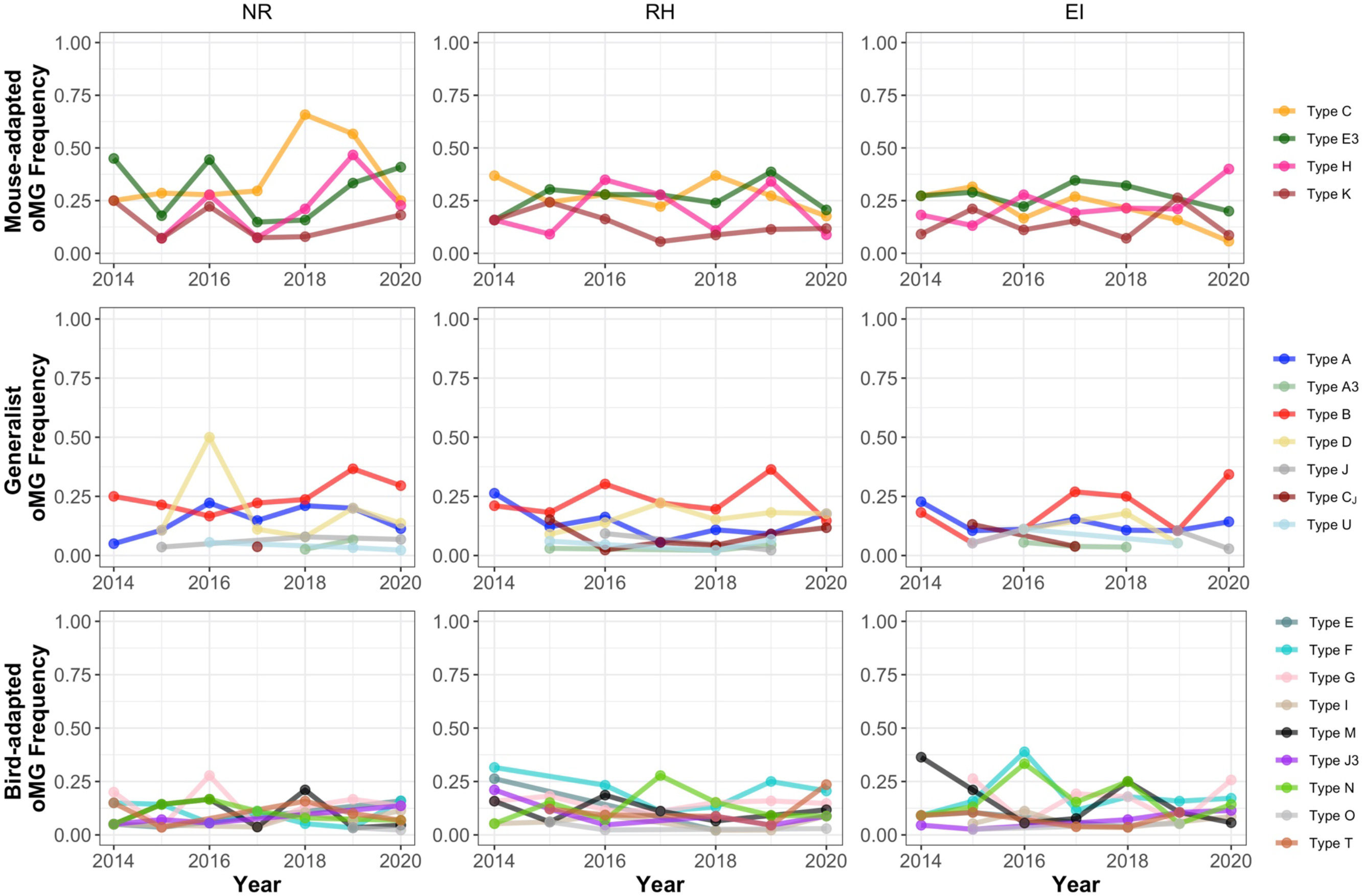
Yearly frequency of individual genotypes in nymphal ticks across sites.

**Figure S6.**
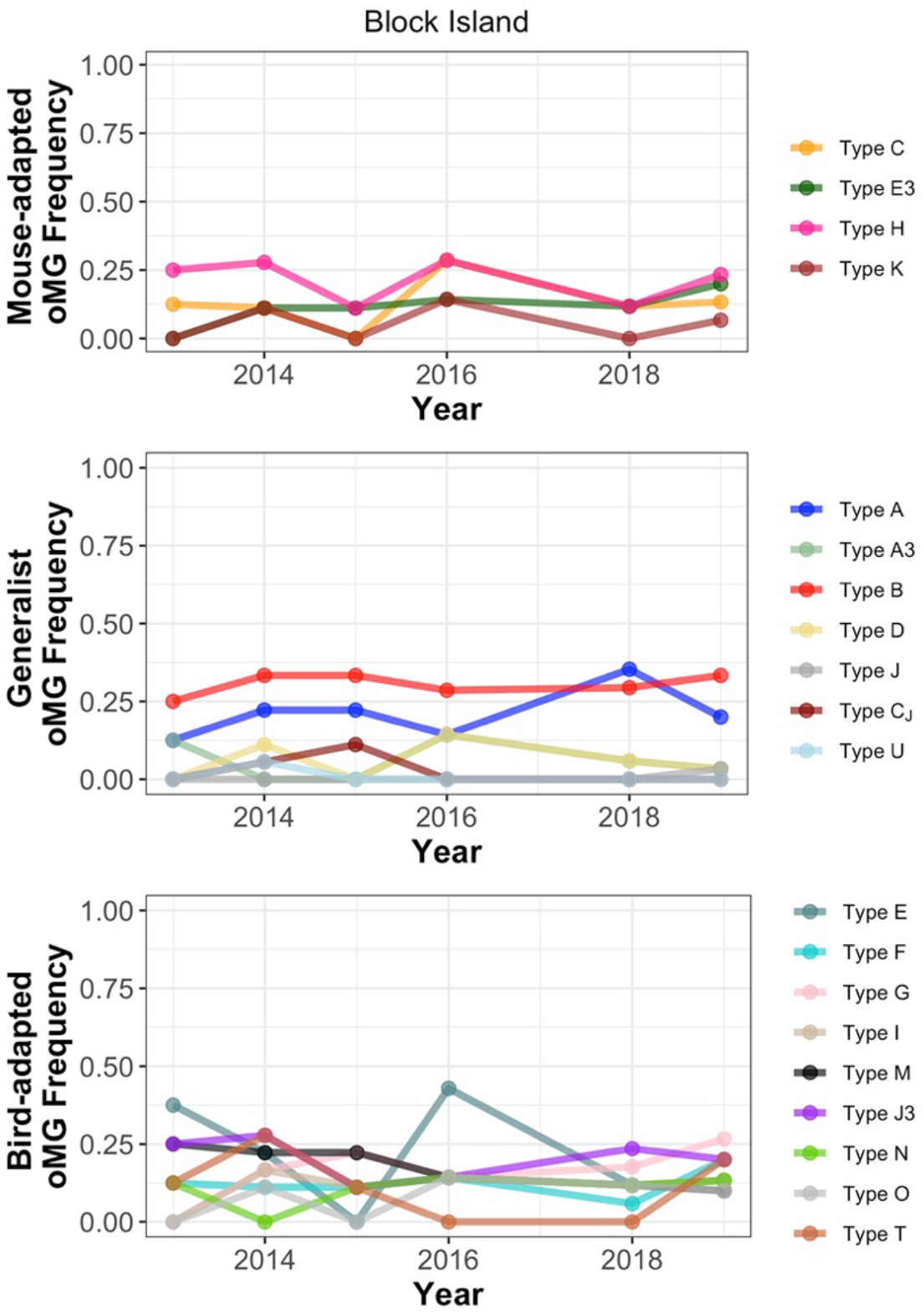
Yearly frequency of individual genotypes in bird hosts, excluding 2017 where n = 2.

**Figure S7.**
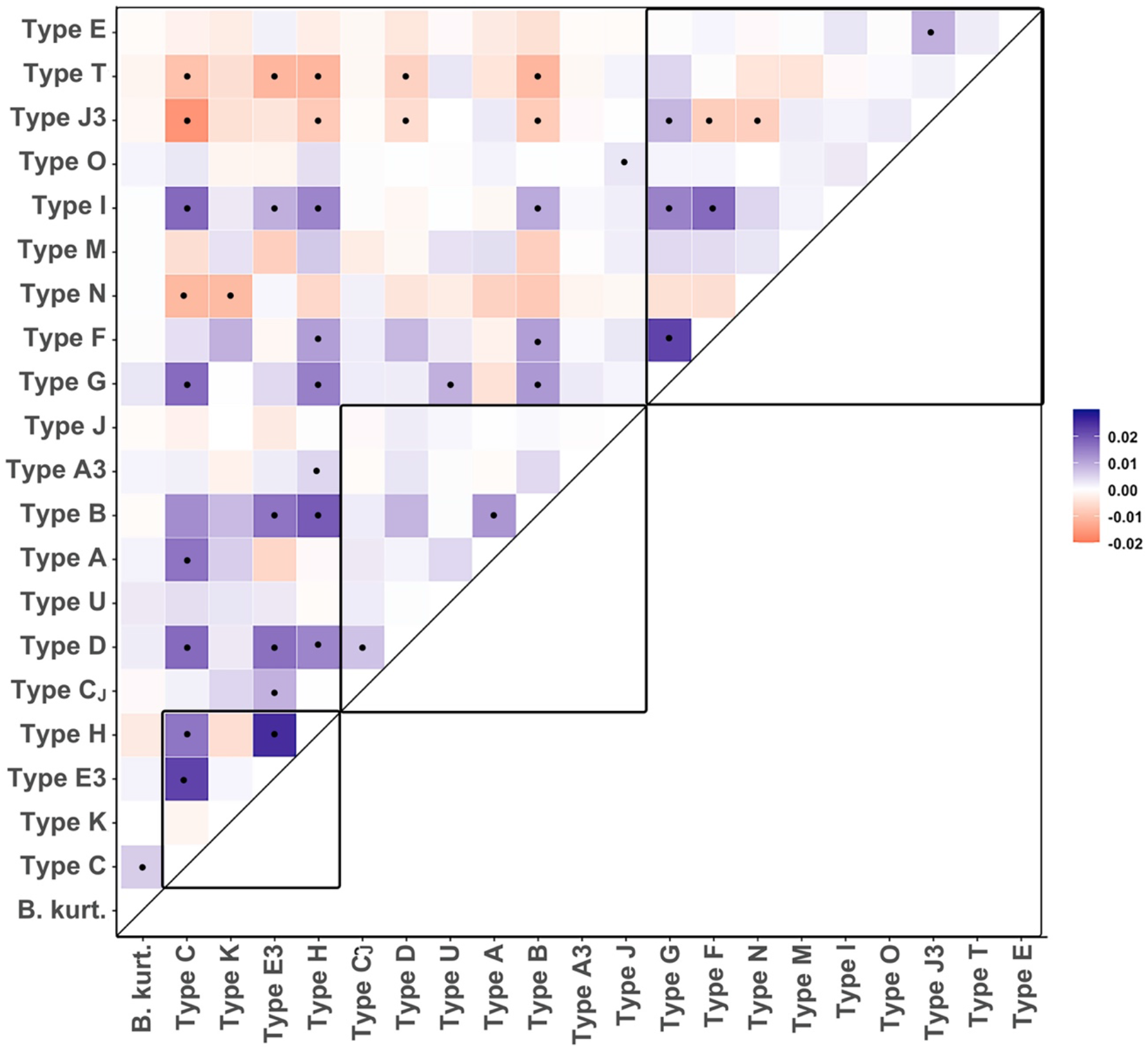
Heatmap of genotype co-occurrence probability effect sizes for nymphal ticks. Color indicates the strength of the effect (blue = positive, red = negative), and black dots indicate significant associations (p < 0.05). Black boxes within the heatmap surround the putative mouse-adapted genotypes (bottom), generalist genotypes (middle), and bird-adapted genotypes (top).

**Figure S8.**
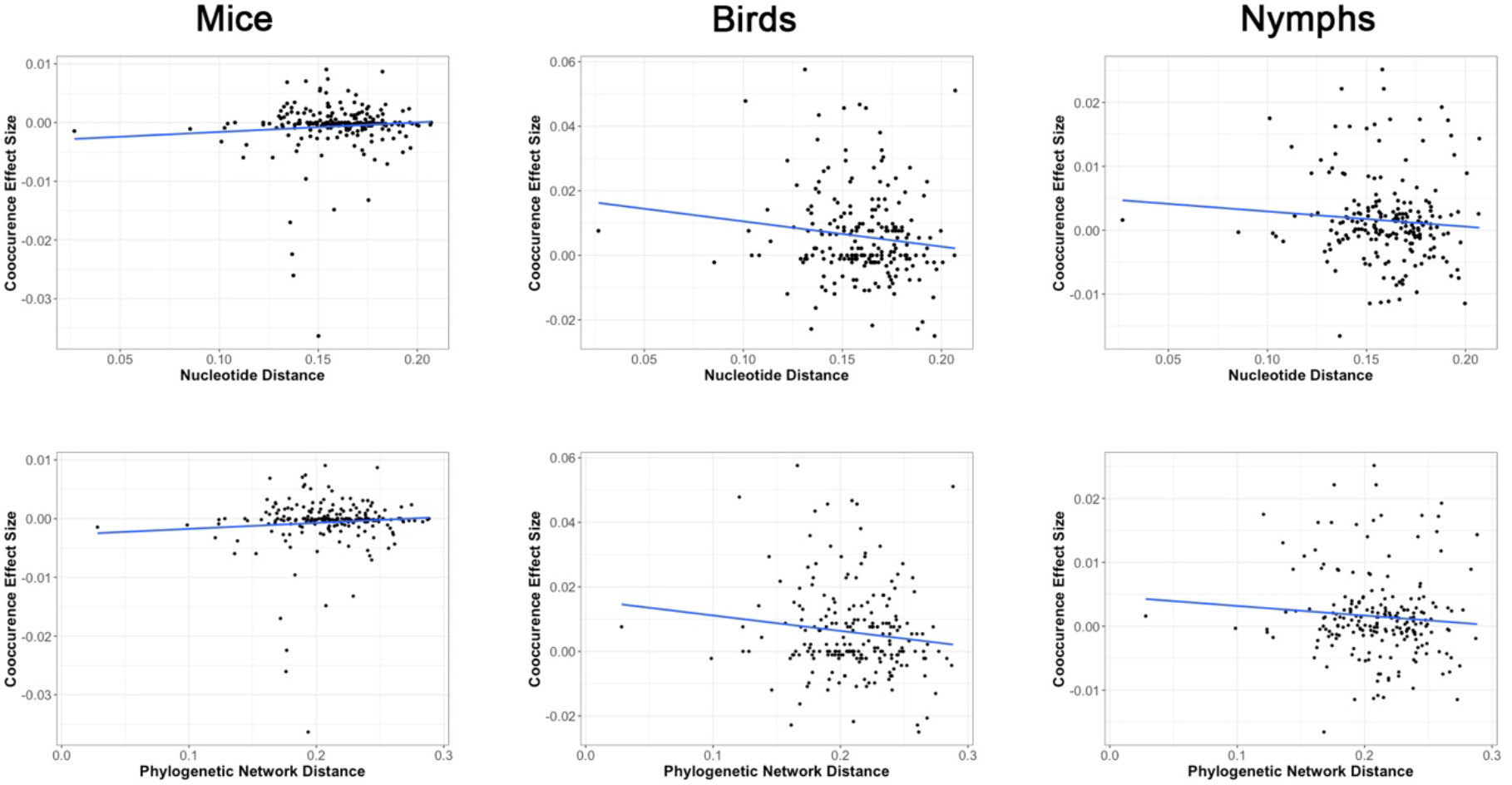
Correlations between pairwise co-occurrence effect sizes among genotypes and nucleotide distance (top row) or phylogenetic network distance (bottom row) among mice (left column), birds (middle column), and nymphal ticks (right column).

**Figure S9.**
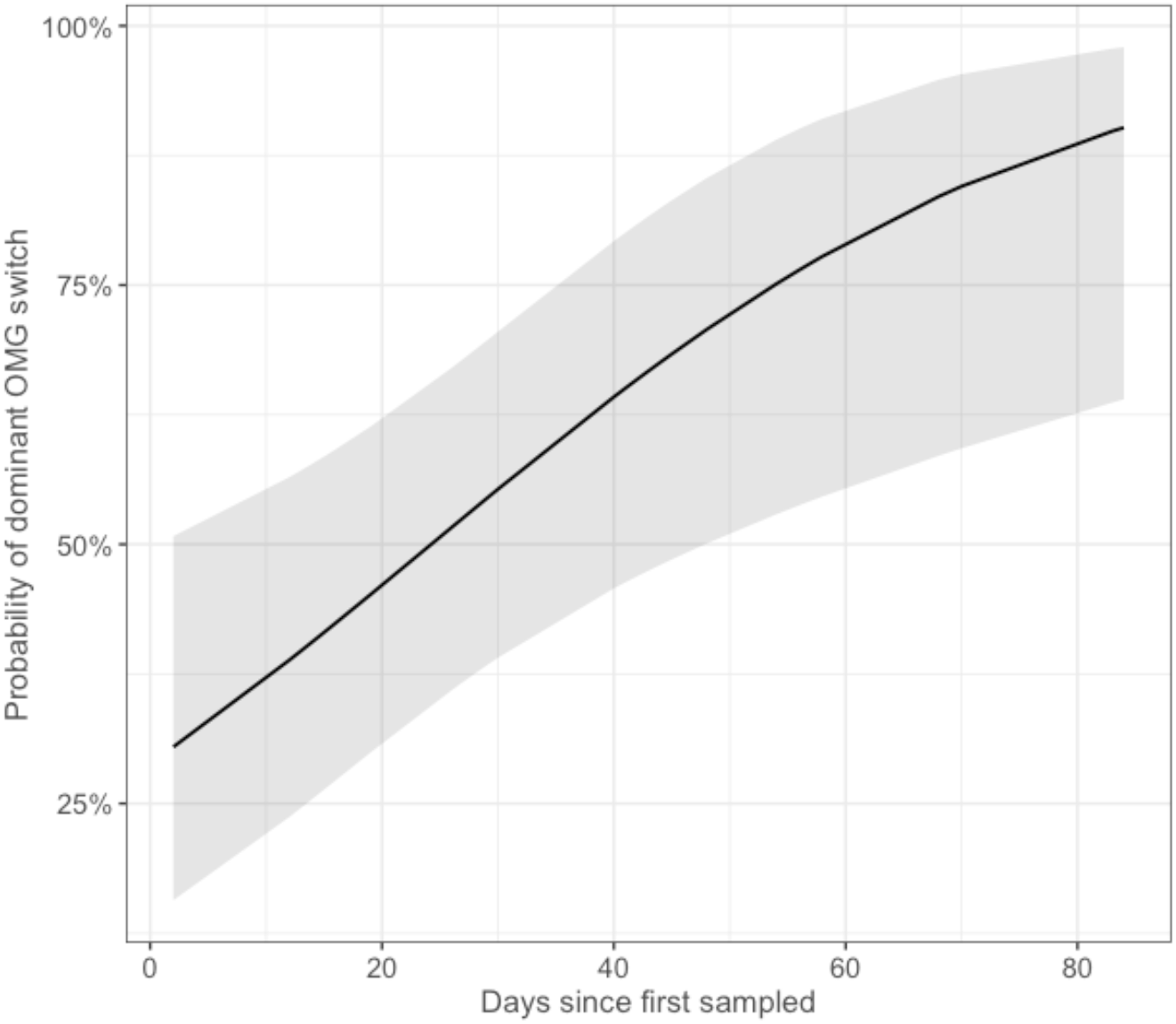
Predicted relationship between time (days) and the probability of turnover in the dominant genotype within individual mammal communities.

**Table S1.** Sequencing of genotypes across *Bb* infected tick nymphs, white footed mice, and passerine birds. Details for individual bird species are provided in Table S3.

| Source | Nymphs |  | Mice |  | Birds |  |
| --- | --- | --- | --- | --- | --- | --- |
| n | 553 |  | 628 |  | 92 |  |
| Richness | 2.6 |  | 1.6 |  | 2.6 |  |
| oMG | Count | Freq. | Count | Freq. | Count | Freq. |
| A | 88 | 0.140 | 66 | 0.119 | 20 | 0.217 |
| B | 146 | 0.232 | 94 | 0.170 | 28 | 0.304 |
| C | 184 | 0.293 | 178 | 0.322 | 12 | 0.130 |
| Cj | 24 | 0.038 | 18 | 0.033 | 2 | 0.022 |
| D | 72 | 0.115 | 46 | 0.083 | 6 | 0.065 |
| E | 22 | 0.035 | 1 | 0.002 | 16 | 0.174 |
| F | 95 | 0.151 | 25 | 0.045 | 14 | 0.152 |
| G | 93 | 0.148 | 37 | 0.067 | 18 | 0.196 |
| H | 133 | 0.212 | 162 | 0.293 | 20 | 0.217 |
| I | 34 | 0.054 | 15 | 0.027 | 12 | 0.130 |
| J | 15 | 0.024 | 2 | 0.004 | 1 | 0.011 |
| K | 82 | 0.131 | 113 | 0.204 | 6 | 0.065 |
| M | 76 | 0.121 | 20 | 0.036 | 14 | 0.152 |
| N | 75 | 0.119 | 16 | 0.029 | 10 | 0.109 |
| O | 8 | 0.013 | 4 | 0.007 | 8 | 0.087 |
| T | 48 | 0.076 | 1 | 0.002 | 14 | 0.152 |
| U | 16 | 0.025 | 4 | 0.007 | 1 | 0.011 |
| A3 | 10 | 0.016 | 5 | 0.009 | 4 | 0.043 |
| E3 | 171 | 0.272 | 144 | 0.260 | 13 | 0.141 |
| J3 | 39 | 0.062 | 2 | 0.004 | 19 | 0.207 |
| B.kurt. | 15 | 0.024 | 27 | 0.049 | 0 | 0.000 |

**Table S2.** Pairwise genotype community similarity among populations of mammals, birds, and nymphs calculated with the local (Sørenson-type) species overlap measure.

|  | Mammals | Birds | Nymphs |
| --- | --- | --- | --- |
| Mammals | 1.000 | 0.672 | 0.904 |
| Birds |  | 1.000 | 0.883 |
| Nymphs |  |  | 1.000 |

**Table S3.**
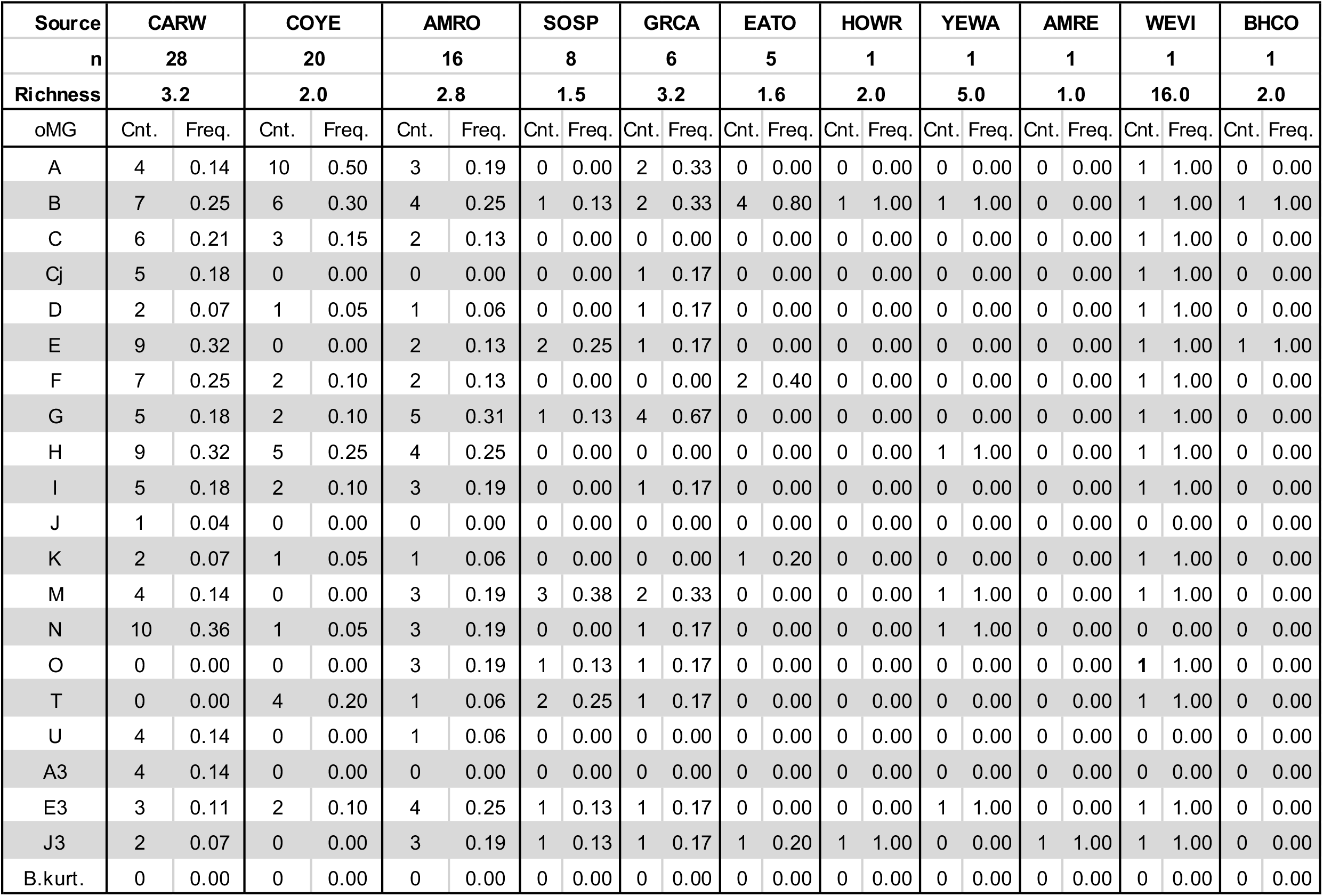
Sequencing of genotypes across individual species of *Bb* passerine birds.

**Table S4.** Mean sojourn time for each infection state as determined by the MSM model. Values represent the expected duration of a mouse infection in days. 95% confidence intervals are provided in parentheses.

|  | Mean Sojourn Time |
| --- | --- |
| Uninfected | 53.9 (41.3 - 71.2) |
| Adapted | 27.6 (19.6 - 38.7) |
| Non-Adapted | 9.5 (6.2 - 14.5) |

**Table S5.**
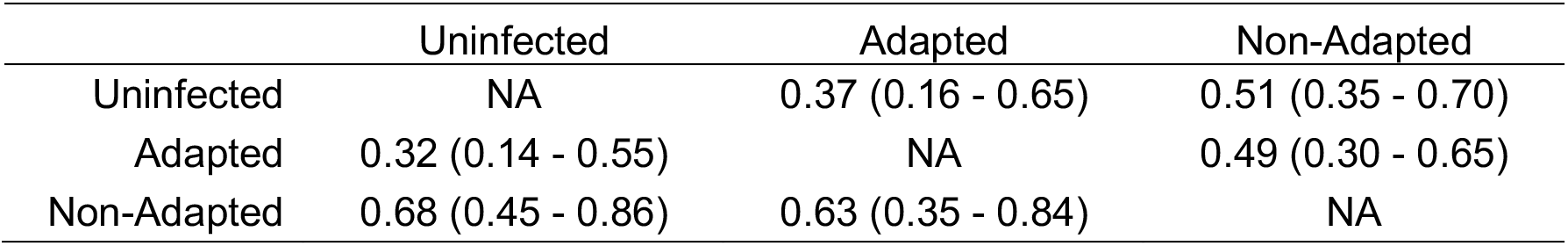
Estimated probabilities of a mouses next different infection state (rows), given the current state (columns), determined by the MSM model. 95% confidence intervals are provided in parentheses.

**Table S6.** Model results for the GLMM testing influence of time and individual mouse characteristics on the probability of observing a switch in the dominant genotype.

| Predictors | Odds Ratios | CI | p |
| --- | --- | --- | --- |
| (Intercept) | 0.51 | 0.20 – 1.30 | 0.164 |
| Time since first sample (days) | 1.03 | 1.01 – 1.06 | **0.006** |
| oMG richness | 1.13 | 0.84 – 1.53 | 0.43 |
| Sex (Male) | 0.92 | 0.50 – 1.71 | 0.796 |
| Age (non-adult) | 1.52 | 0.28 – 11.41 | 0.643 |
| Nmphal Burden | 0.99 | 0.94 – 1.06 | 0.862 |

**Table S7.** Recombination analysis results.

| In Alignment |  | In Recombinant Sequence |  |  |  |  | Detection Method |  |  |  |  |  |  |
| --- | --- | --- | --- | --- | --- | --- | --- | --- | --- | --- | --- | --- | --- |
| Begin | End | Begin | End | Recombinant Sequence(s) | Minor Parental Sequence(s) | Major Parental Sequence(s) | RDP | GENECONV | Bootscan | Maxchi | Chimaera | SiScan | 3Seq |
| 303 | 413 | 297 | 398 | B | Cj, C | J, H | 5.15E-05 | 2.21E-04 | 2.18E-06 | 1.62E-07 | 1.37E-07 | 1.62E-09 | 1.12E-07 |
| 459 | 554 | 441 | 530 | B.kurt | J | C | NS | 2.84E-07 | 1.28E-08 | 2.56E-04 | 2.56E-04 | 1.63E-05 | 8.66E-08 |
| 350 | 408 | 335 | 393 | I | D | Cj, C | 3.84E-07 | 2.78E-06 | 1.53E-06 | 4.60E-03 | 7.96E-04 | 2.64E-05 | 2.38E-05 |
| 303 | 493 | 297 | 469 | J3 | A | M | NS | 2.38E-03 | 9.61E-04 | 5.91E-06 | 9.63E-03 | 3.89E-07 | 6.14E-07 |
| 142 | 418 | 139 | 403 | A | G | C, Cj | NS | NS | NS | 5.17E-05 | 4.68E-06 | 4.50E-03 | 4.55E-04 |
| 320 | 526 | 317 | 508 | B.kurt | E | H | NS | NS | NS | 1.91E-05 | 4.38E-02 | NS | 1.03E-02 |
| 628 | 186 | 604 | 183 | H | O | B.kurt | NS | 2.22E-03 | 1.96E-03 | 1.88E-02 | 1.43E-02 | NS | 2.22E-03 |
| 518 | 198 | 497 | 195 | E3 | U | G | NS | NS | 8.77E-03 | 2.09E-04 | 5.25E-05 | 4.67E-04 | NS |
| 608 | 294 | 587 | 291 | K | J | N | NS | NS | 8.03E-04 | NS | 5.73E-03 | NS | NS |
| 250 | 309 | 247 | 303 | J3 | E | A | NS | NS | NS | 2.51E-03 | NS | 1.49E-03 | 1.66E-03 |
| 340 | 418 | 326 | 400 | O | C, Cj | J, H | NS | NS | 2.19E-03 | 4.52E-03 | 1.66E-03 | 2.85E-05 | 4.28E-02 |

**Table S8.** Accession numbers for strains used for alignment during primer design.

| Accession | Strain |
| --- | --- |
| **CP001550** | 29805 |
| **CP002239** | N40 |
| **CP001446** | WI91-23 |
| **CP017202** | B331 |
| **CP001535** | 118a |
| **CP001375** | 72a |
| **CP001484** | CA-11.2A |
| **CP001493** | 94a |
| **CP031398** | MM1 |
| **CP002316** | JD1 |
| **CP001271** | 156a |
| **CP002268** | 297 |
| **CP001422** | 64b |
| **CP001212** | ZS7 |
| **AE000792** | B31 |
| **CP019917** | Pabe |
| **CP019755** | B31_NRZ |
| **CP001568** | Bol26 |

**Table S9.**
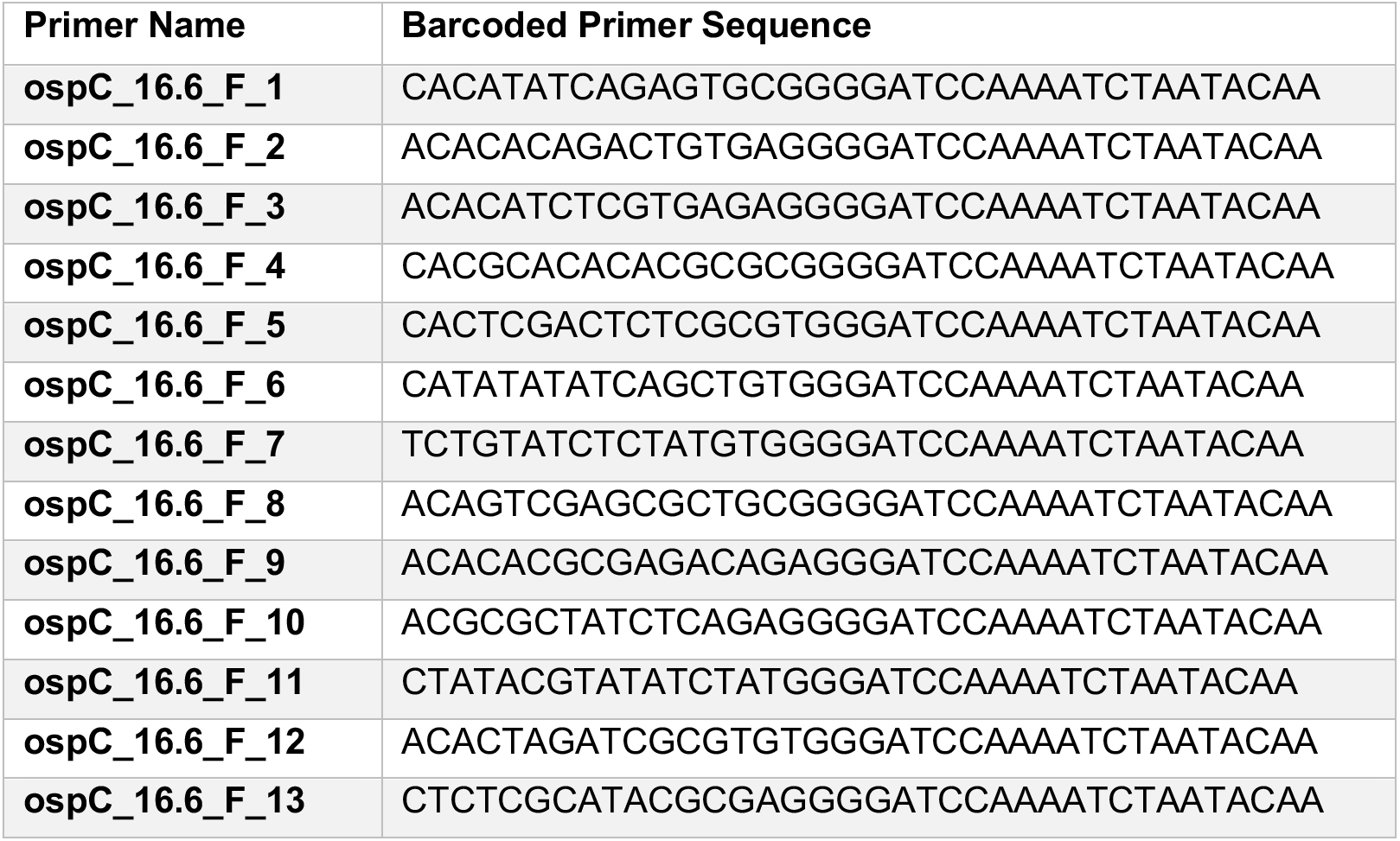
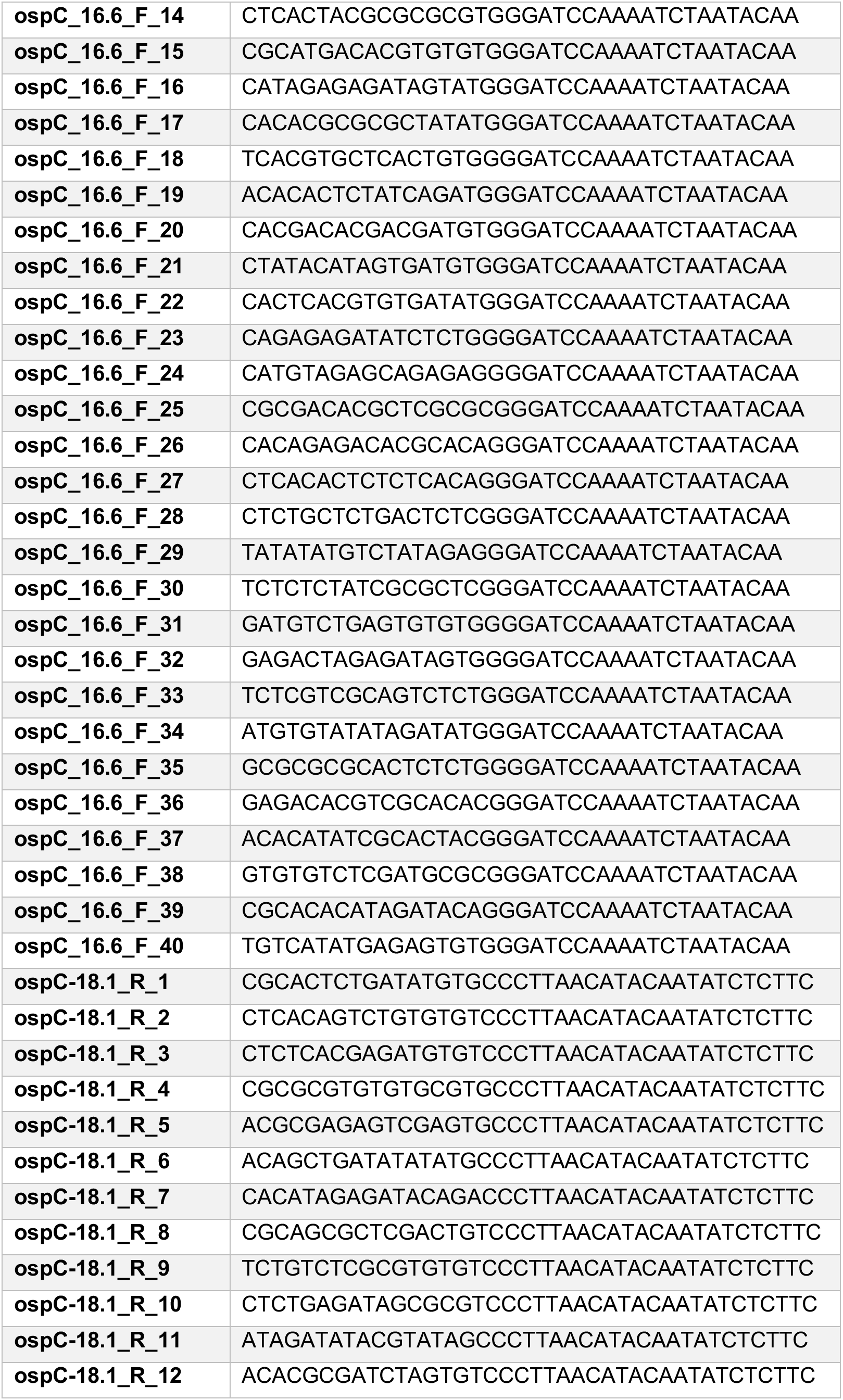
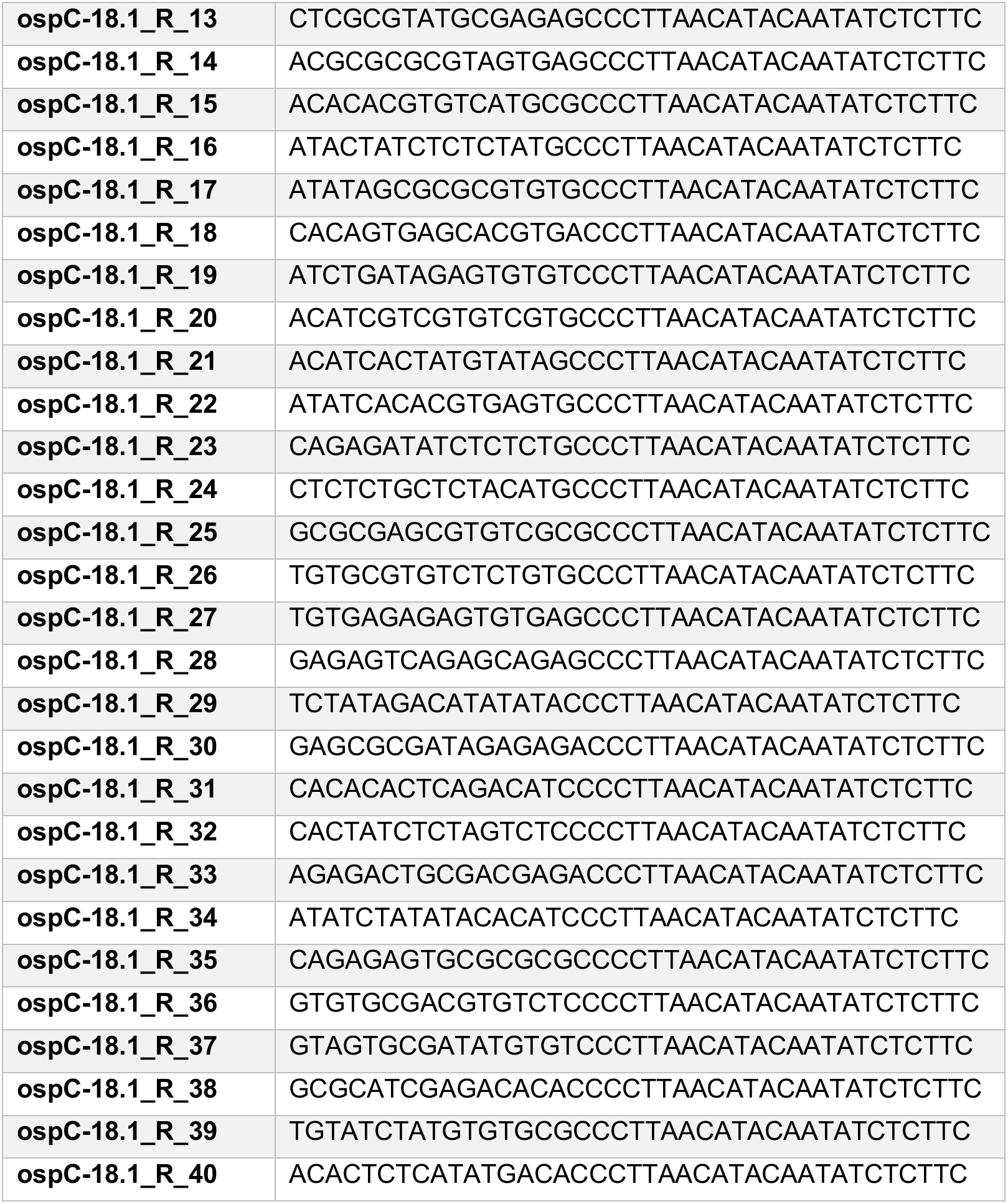
Barcoded *ospC* primer set 1 used in this study to amplify 1500bp region centered around the ospC locus. Barcodes selected from a set of 384 sequences provided by Pacific <u>Biosciences.</u>

**Table S10.**
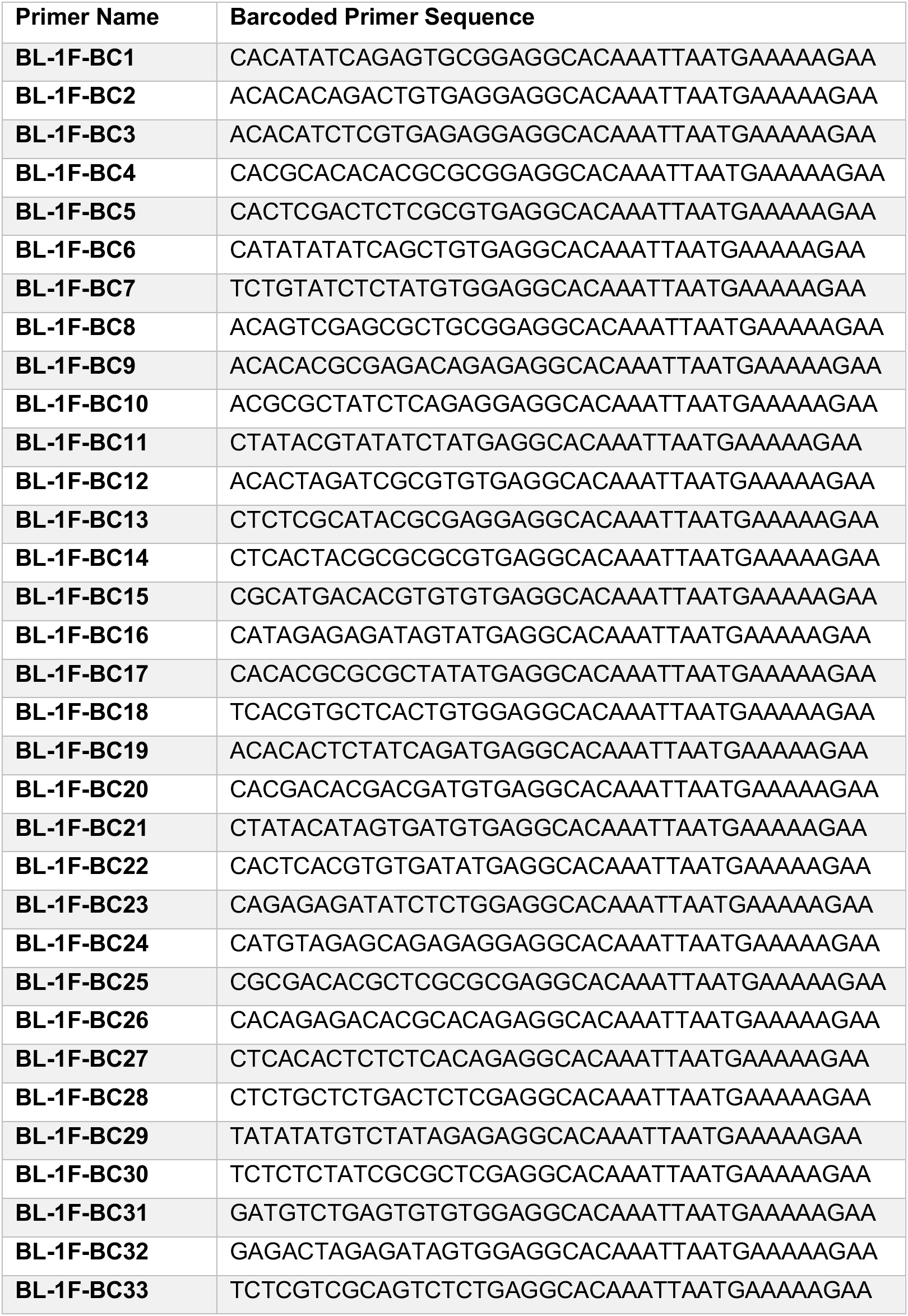
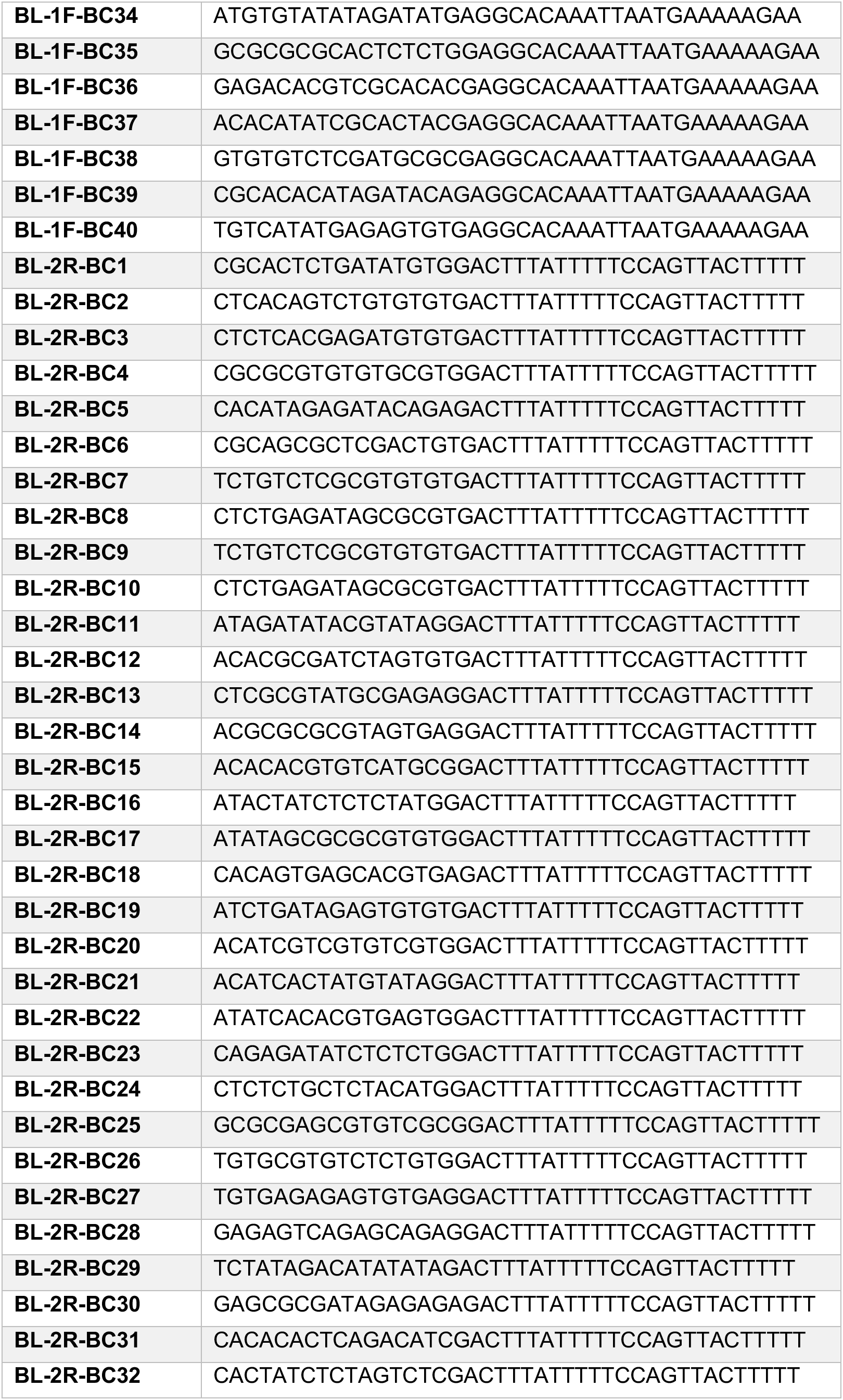
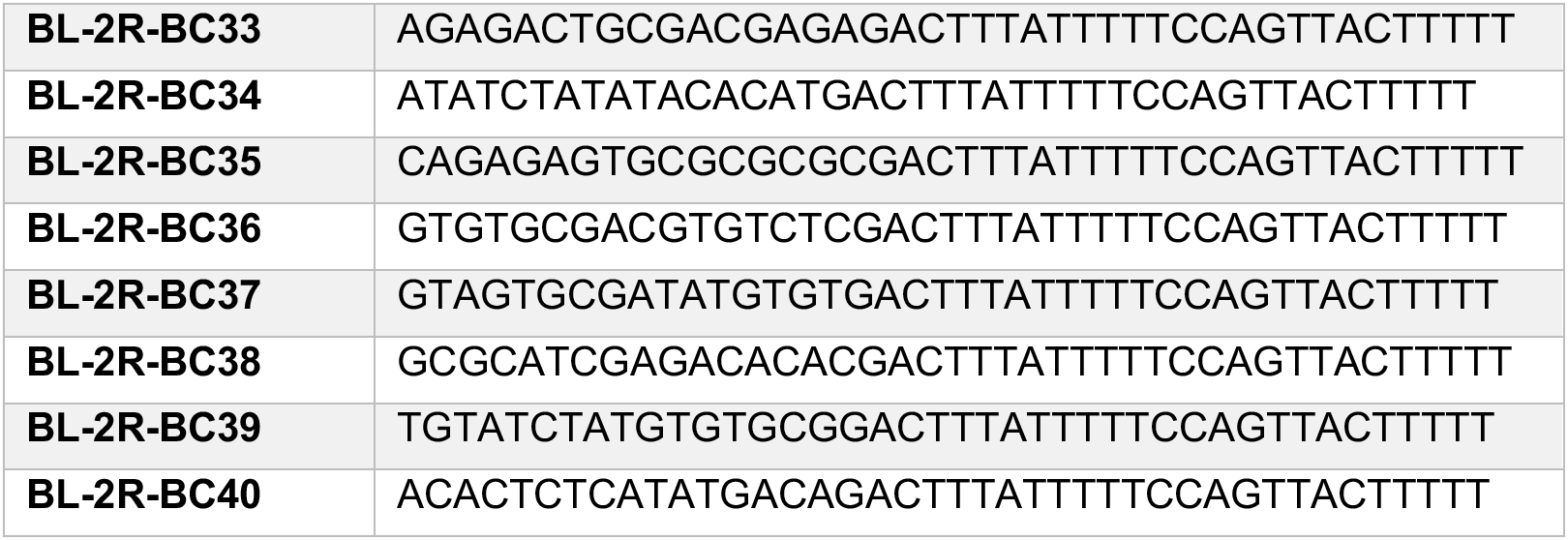
Barcoded *ospC* primer set 2 used in this study to amplify 750bp region encompassing the *ospC* locus. Barcodes selected from a set of 384 sequences provided by Pacific Biosciences.

**Table S11.**
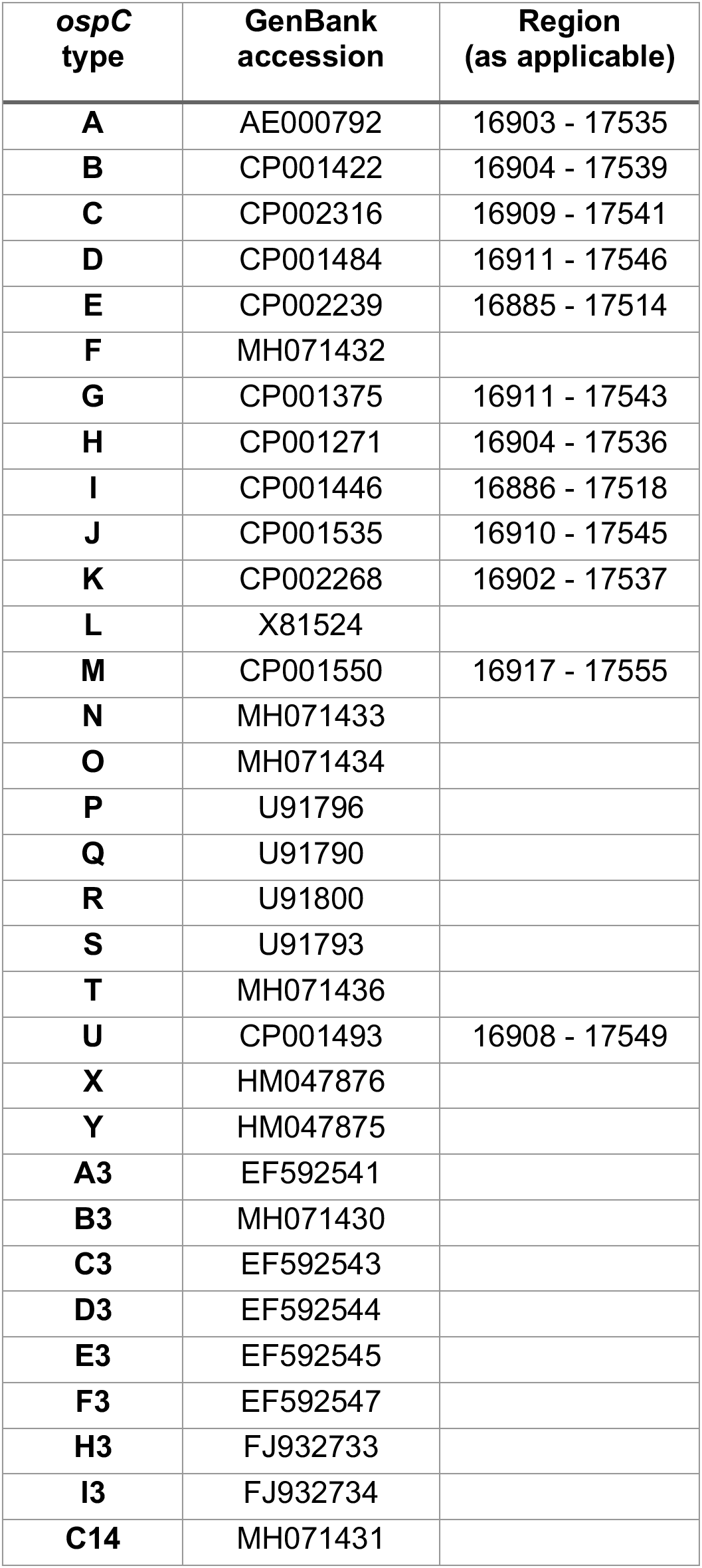
Accession numbers of reference *ospC* sequences used during sequence clustering.

| ***ospC***<br>**type** | **GenBank**<br>**accession** | **Region**<br>**(as applicable)** |
| --- | --- | --- |
| **A** | AE000792 | 16903 - 17535 |
| **B** | CP001422 | 16904 - 17539 |
| **C** | CP002316 | 16909 - 17541 |
| **D** | CP001484 | 16911 - 17546 |
| **E** | CP002239 | 16885 - 17514 |
| **F** | MH071432 |  |
| **G** | CP001375 | 16911 - 17543 |
| **H** | CP001271 | 16904 - 17536 |
| **I** | CP001446 | 16886 - 17518 |
| **J** | CP001535 | 16910 - 17545 |
| **K** | CP002268 | 16902 - 17537 |
| **L** | X81524 |  |
| **M** | CP001550 | 16917 - 17555 |
| **N** | MH071433 |  |
| **O** | MH071434 |  |
| **P** | U91796 |  |
| **Q** | U91790 |  |
| **R** | U91800 |  |
| **S** | U91793 |  |
| **T** | MH071436 |  |
| **U** | CP001493 | 16908 - 17549 |
| **X** | HM047876 |  |
| **Y** | HM047875 |  |
| **A3** | EF592541 |  |
| **B3** | MH071430 |  |
| **C3** | EF592543 |  |
| **D3** | EF592544 |  |
| **E3** | EF592545 |  |
| **F3** | EF592547 |  |
| **H3** | FJ932733 |  |
| **I3** | FJ932734 |  |
| **C14** | MH071431 |  |

